# Off-target-free chemogenetic platform that decodes physiological roles of target GPCRs

**DOI:** 10.64898/2026.05.24.727545

**Authors:** Yuma Matsuoka, Hajime Inoue, Tatsuki Hori, Duy Phuoc Tran, Akio Kitao, Tomohiro Doura, Shigeki Kiyonaka

## Abstract

G-protein-coupled receptors (GPCRs) are essential mediators of cellular processes, and their dysfunction is implicated in various diseases. Although pharmacological approaches are powerful for elucidating GPCR functions, careful consideration of potential off-target effects remains crucial. Here, we present an off-target-free chemogenetic platform to dissect the physiological roles of target GPCRs. As a proof-of-concept, we focused on the adenosine A_2A_ receptor (A_2A_R), a prototypical GPCR involved in neurodegenerative diseases. Leveraging the ligand recognition mechanisms of A_2A_R, we designed a prominent chemogenetic pair comprising a clinical A_2A_R antagonist and an engineered A_2A_R mutant that is insensitive to the antagonist but retains responsiveness to adenosine. Using this pair, we uncovered the critical role of A_2A_R in neurite outgrowth under hypoxic conditions in neuron-like cells, while avoiding potential off-target effects. Furthermore, we demonstrate the generalizability of this chemogenetic approach to other class A GPCRs, offering a versatile platform for receptor-specific dissection of GPCR signaling.

## INTRODUCTION

Precise control of proteins of interest (POIs) in living cells is essential for understanding the physiological roles of POIs. For this purpose, genetic or pharmacological methods are widely applied. Genetic approaches, such as knocking out POIs, enable loss-of-function analyses under cellular conditions.^1^ Although genetic methods offer high target selectivity, these approaches are less suitable for investigating POI functions over short timeframes. In contrast, pharmacological approaches using selective inhibitors enable acute and reversible inhibition of POI functions. Despite their advantages, the use of inhibitors often carries the risk of off-target effects.^2,3^ Therefore, even with highly selective ligands, careful consideration of unforeseen off-target effects is essential.

G-protein-coupled receptors (GPCRs) represent the largest class of cell surface receptors.^4^ GPCRs activate downstream signaling pathways that play key roles in various physiological processes. Dysregulation of these receptors is directly linked to various diseases, making GPCRs important targets for drug development. Notably, approximately 30% of drugs approved by the US Food and Drug Administration (FDA) target GPCRs, and selective inhibitors for various GPCRs have been developed.^5^ Although these compounds are powerful for elucidating the physiological roles of target GPCRs, off-target effects resulting from unintended binding or metabolic byproducts are frequently observed. Thus, a new approach that offers a precise evaluation of target GPCR signaling pathways with minimal off-target effects is highly desirable.

Adenosine receptors are a family of GPCRs activated by the endogenous ligand adenosine, a product primarily derived from the metabolism of adenosine triphosphate (ATP).^6^ A_2A_R is one of four known adenosine receptors (A_1_R, A_2A_R, A_2B_R, and A_3_R), and is recognized as a prototypical GPCR that is highly expressed in the nervous system and immune cells.^7^ A_2A_R plays essential roles in regulating the sleep‒wake cycle and motor activity in the brain,^8,9^ and also contributes to immune function and angiogenesis in response to adenosine.^10,11^ Dysregulation of A_2A_R has been implicated in neurodegenerative diseases such as Parkinson’s disease and Alzheimer’s disease,^12,13^ as well as cancer.^10^ Therefore, A_2A_R has attracted considerable attention as a potential drug target, and many A_2A_R-selective antagonists have been developed, with some currently in clinical trials. These compounds are also beneficial for elucidating the physiological roles of A_2A_R in living cells. However, given the potential for off-target binding or the formation of metabolic byproducts, it remains challenging to demonstrate that observed drug effects are purely mediated by A_2A_R and its downstream signaling.

In this report, we present an off-target-free chemogenetic strategy for elucidating the physiological roles of target GPCRs in living cells. Conventionally, receptor chemogenetics has been used to activate receptor signaling in a cell-type-specific manner (for details, see DISCUSSION). Here, we applied chemogenetics to clarify the physiological roles of target GPCRs. For this purpose, we focused on A_2A_R as a model target and designed A_2A_R mutants that are insensitive to a clinically validated A_2A_R antagonist while retaining full responsiveness to adenosine. We also demonstrated that this chemogenetic approach is applicable to a serotonin receptor, a different class A GPCR family member. Furthermore, using a chemogenetic pair consisting of an engineered A_2A_R and its selective antagonist, we revealed the physiological role of A_2A_R under hypoxic conditions in neuron-like cells.

## Results

### Design of antagonist-insensitive A_2A_R mutants that retain adenosine responsiveness

To investigate the physiological roles of A_2A_R, we reasoned that antagonist-insensitive A_2A_R mutants that retain responsiveness to adenosine would serve as powerful tools for this purpose. In systems where A_2A_R activation drives physiological responses such as neuronal differentiation, these responses can be suppressed by A_2A_R-selective antagonists (**Figure 1A**, left). However, under these conditions, the contribution of unforeseen off-target effects cannot be excluded. In contrast, when antagonists are applied to cells expressing an antagonist-insensitive A_2A_R mutant, the cellular responses are induced exclusively through off-target effects (**Figure 1A**, right). Therefore, comparing wild-type (WT) A_2A_R with an antagonist-insensitive mutant enables the analysis of the physiological roles of A_2A_R by eliminating off-target effects. However, given that antagonists share the same binding site as endogenous agonists, designing antagonist-insensitive A_2A_R mutants that retain adenosine responsiveness remains highly challenging.

**Figure 1.**
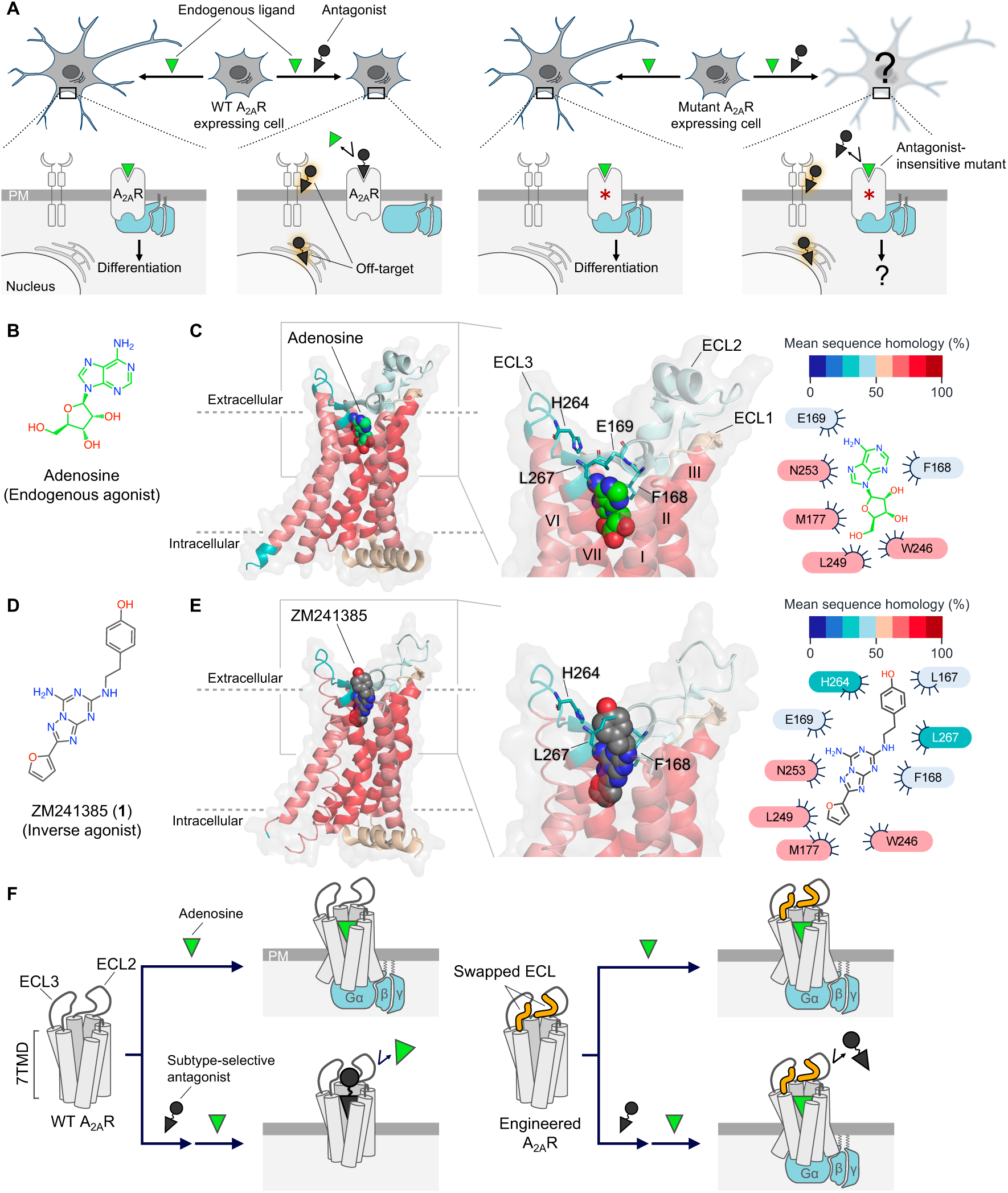
Design of antagonist-insensitive A_2A_R mutants retaining adenosine responsiveness. (**A**) Left, schematic illustration of on-target and off-target effects by antagonists. Right, strategy for assessing off-target effects using antagonist-insensitive mutants. PM, plasma membrane. (**B**) Chemical structure of adenosine. (**C**) X-ray structure of A_2A_R bound to adenosine (Protein Data Bank (PDB): 2YDO). Left and middle, receptor structure coloured according to mean sequence homology among adenosine receptor subtypes. Right, two-dimensional diagram of receptor residues interacting with adenosine. (**D**) Chemical structure of ZM241385, an A_2A_R-selective inverse agonist. (**E**) X-ray structure of A_2A_R bound to ZM241385 (PDB: 3EML). Left and middle, receptor structure coloured as in (C). Right, two-dimensional diagram of residues interacting with ZM241385. (**F**) Overview of engineered A_2A_R mutants designed to prevent the antagonist binding while retaining adenosine responsiveness. Mutated residues in extracellular loops (ECLs) are highlighted in yellow.

We focused on the ligand recognition mechanisms of A_2A_R to design antagonist-insensitive mutants. A_2A_R recognizes both adenosine and antagonists within the seven transmembrane domains (7TMD) (**Figure 1, B** to **E**).^14,15^ Sequence homology among adenosine receptors is shown as a heatmap in **Figures 1C** and **1E**. Although the 7TMD, where adenosine primarily binds, is highly conserved among adenosine receptor subtypes, we found a markedly lower sequence homology in the extracellular loops (ECLs) (**Figures 1C** and **S1**). A_2A_R-selective antagonists were anticipated to achieve high A_2A_ subtype selectivity by interacting with both the ECLs and the 7TMD. In the case of ZM241385 (**1**), a well-known A_2A_R-selective antagonist, the X-ray structure revealed that the phenol group, which extends toward the extracellular surface of A_2A_R, interacts with Leu167 on ECL2 and His264 and Leu267 on ECL3 (**Figure 1E**). Notably, these residues are less conserved among other adenosine receptors (**Figures 1E** and **S1**). Therefore, we hypothesized that the engineered A_2A_R mutants, in which parts of the ECLs of A_2A_R are swapped with those from other adenosine receptors, can evade the inhibitory effects of A_2A_R-selective antagonists. Of note, introducing mutations in the ECLs that are spatially distant from the adenosine binding site in the 7TMD should enable the generation of antagonist-insensitive A_2A_R mutants that retain adenosine responsiveness. As shown in **Figure 1F**, the engineered A_2A_R is expected to remain responsive to endogenously released adenosine, even in the presence of an antagonist.

### Functional assay of engineered A_2A_Rs using adenosine and ZM241385

In designing mutations to exchange ECLs in A_2A_R, we focused on ECL2 and ECL3, both of which are located above the ligand-binding domain of A_2A_R. Here, three A_2A_R mutants, A_2A_R(ECL-A_1_), A_2A_R(ECL-A_2B_), and A_2A_R(ECL-A_3_), were constructed by swapping Leu167–Val172 of ECL2 and Pro260–Leu267 of ECL3 in A_2A_R with the corresponding residues from A_1_R, A_2B_R, and A_3_R (**Figure 2A**). Notably, Leu167, His264, and Leu267, which potentially contribute to subtype selectivity of ZM241385, are included in the mutation sites.

**Figure 2.**
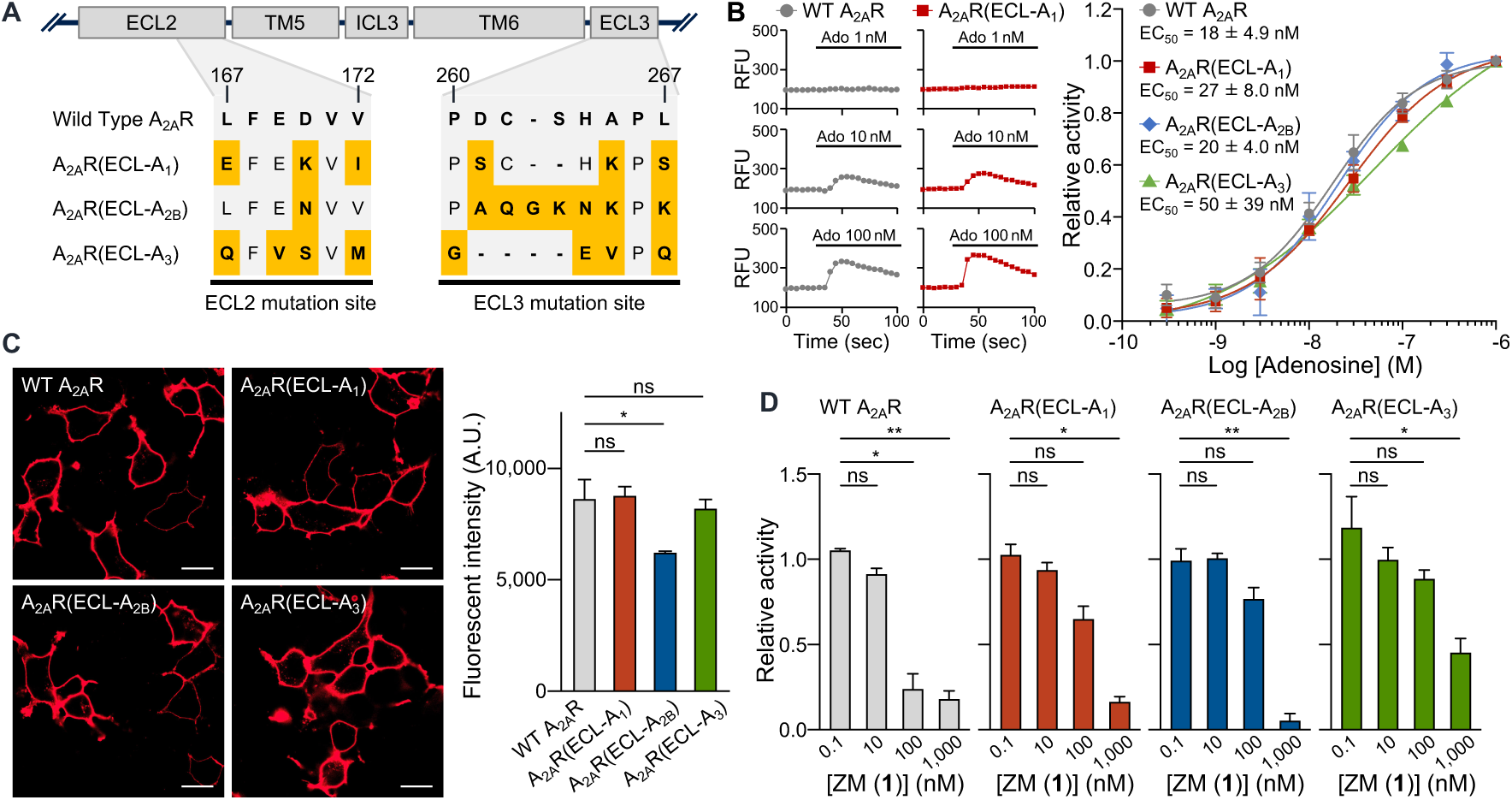
Construction and characterization of engineered A_2A_R chimeras. (**A**) Mutation sites of chimeric A_2A_R mutants. Residues differing from WT A_2A_R are highlighted in yellow. (**B**) Adenosine-induced Ca^2+^ responses in HEK293 cells co-expressing Gα_15_. Left, representative traces for WT A_2A_R (gray) and A_2A_R(ECL-A_1_) (red) stimulated by adenosine (Ado, black bar). Right, Concentration–response curves for cells expressing WT A_2A_R (gray), A_2A_R(ECL-A_1_) (red), A_2A_R(ECL-A_2B_) (blue), or A_2A_R(ECL-A_3_) (green). Relative activity is normalized to ΔRFU at 1 µM Ado. *N* = 3 independent experiments. (**C**) Confocal images of A_2A_Rs in HEK293 cells. Left, representative images of HEK293 cells expressing 2×HA-tagged WT A_2A_R or chimeric A_2A_R mutants, stained with Alexa 647-conjugated anti-HA tag antibody. Scale bar, 20 µm. Right, quantification of cell-surface fluorescent intensity. *n* = 3 independent experiments, 10 cells per condition. One-way ANOVA with Dunnett’s test versus WT A_2A_R. (**D**) Inhibition of 100 nM Ado-induced Ca^2+^ responses by ZM241385 (ZM). Relative activity is normalized to ΔRFU measured at 100 nM Ado alone. *n* = 3 independent experiments. One-way ANOVA with Dunnett’s test versus 0.1 nM ZM. Data are mean ± s.e.m. \**P* < 0.05, \*\**P* < 0.01; ns, not significant.

We investigated whether these A_2A_R chimeric mutants retain adenosine responsiveness comparable to that of WT A_2A_R. A_2A_R activity was assessed by monitoring changes in intracellular Ca^2+^ concentrations using a fluorescent Ca^2+^ indicator, Cal-520 AM, after ligand treatment in HEK293 cells transfected with each A_2A_R mutant and Gα_15_ protein.^16^ Treatment with adenosine to these cells increased Cal-520 fluorescence in a dose-dependent manner (**Figure S2**). As expected, both the intensity of Ca^2+^ responses and the calculated half-maximal effective concentration (EC_50_) values for adenosine in the three A_2A_R mutants were comparable to WT A_2A_R (**Figure 2B**). We also investigated the expression levels of these A_2A_R mutants on the plasma membrane by immunostaining under live-cell conditions, using A_2A_R mutants fused with a tandem HA-tag (2×HA) at the N-terminus. As shown in **Figure 2C**, confocal imaging showed that the expression level of A_2A_R(ECL-A_2B_) was slightly reduced compared with WT A_2A_R (**Figures 2C** and **S3**). In contrast, A_2A_R(ECL-A_1_) and A_2A_R(ECL-A_3_) retained their expression levels on the cell surface.

Next, the inhibitory effects of ZM241385 on these mutants were examined in HEK293 cells using the Ca^2+^ mobilization assay. As shown in **Figure S4**, each concentration of ZM241385 was pretreated, and adenosine-induced Ca^2+^ responses were evaluated. Consistent with previous studies,^17^ 100 nM of ZM241385 effectively suppressed adenosine-induced responses in WT A_2A_R (**Figure 2D**). However, the same concentration of ZM241385 was insufficient to suppress the Ca^2+^ responses mediated by the A_2A_R mutants (A_2A_R(ECL-A_1_), A_2A_R(ECL-A_2B_), and A_2A_R(ECL-A_3_)). A higher concentration (1,000 nM) of ZM241385 suppressed the responses of both WT A_2A_R and the mutants. These results demonstrate that replacing the ECLs of A_2A_R with those of other adenosine receptor subtypes weakens the inhibitory effect of the A_2A_R-selective antagonist while maintaining the response to the endogenous ligand, adenosine. Given their preserved responsiveness to adenosine and appropriate expression level on the plasma membrane, A_2A_R(ECL-A_1_) and A_2A_R(ECL-A_3_) were considered potential mutants for our purposes. Here, we selected A_2A_R(ECL-A_1_) in subsequent experiments because Cys262 on ECL3 is conserved, as it contributes to the formation of a disulfide bond with another cysteine residue. ^15^

### Preladenant, a suitable ligand for the antagonist-insensitive A_2A_R system

A_2A_R(ECL-A_1_) successfully reduced the inhibitory effect of ZM241385 compared with WT A_2A_R, without affecting the adenosine-induced response. However, the reduced inhibition was insufficient for practical applications. Antagonists with significantly reduced inhibitory effects against A_2A_R(ECL-A_1_) were identified by conducting inhibitory assays using A_2A_R-selective antagonists with diverse chemical structures, including clinically validated antagonists (**Figure 3A**). Our assay revealed that the inhibitory effects were primarily influenced by the chemical structures of the antagonists (**Figure 3B**). In the case of ZM241385 (**1**), the half-maximal inhibitory concentration (IC_50_) was 4.6-fold higher for A_2A_R(ECL-A_1_) than WT A_2A_R. No prominent changes in IC_50_ values were observed between WT A_2A_R and A_2A_R(ECL-A_1_) for CGS15943 (**2**), imaradenant (**3**), vipadenant (**4**), ciforadenant (**5**), and etrumadenant (**6**). Importantly, IC_50_ values for A_2A_R(ECL-A_1_) were 9.5-fold and over 1,000-fold higher than those of WT A_2A_R with SCH442416 (**7**) and preladenant (**8**), respectively. For several of these antagonists, X-ray structures of A_2A_R in complex with these compounds are available,^18–20^ enabling detailed analysis of structure‒function relationships (**Figure 3C**). In the case of imaradenant (**3**), vipadenant (**4**), and etrumadenant (**6**), these antagonists primarily interact with 7TMD. In contrast, preladenant (**8**) interacts with 7TMD and ECLs. In particular, the phenylpiperazinylethyl moiety attached to N7 in preladenant (**8**) interacts with the extracellular surface of A_2A_R, specifically stabilized by π‒π stacking with His264 on ECL3 (**Figure 3C**).

**Figure 3.**
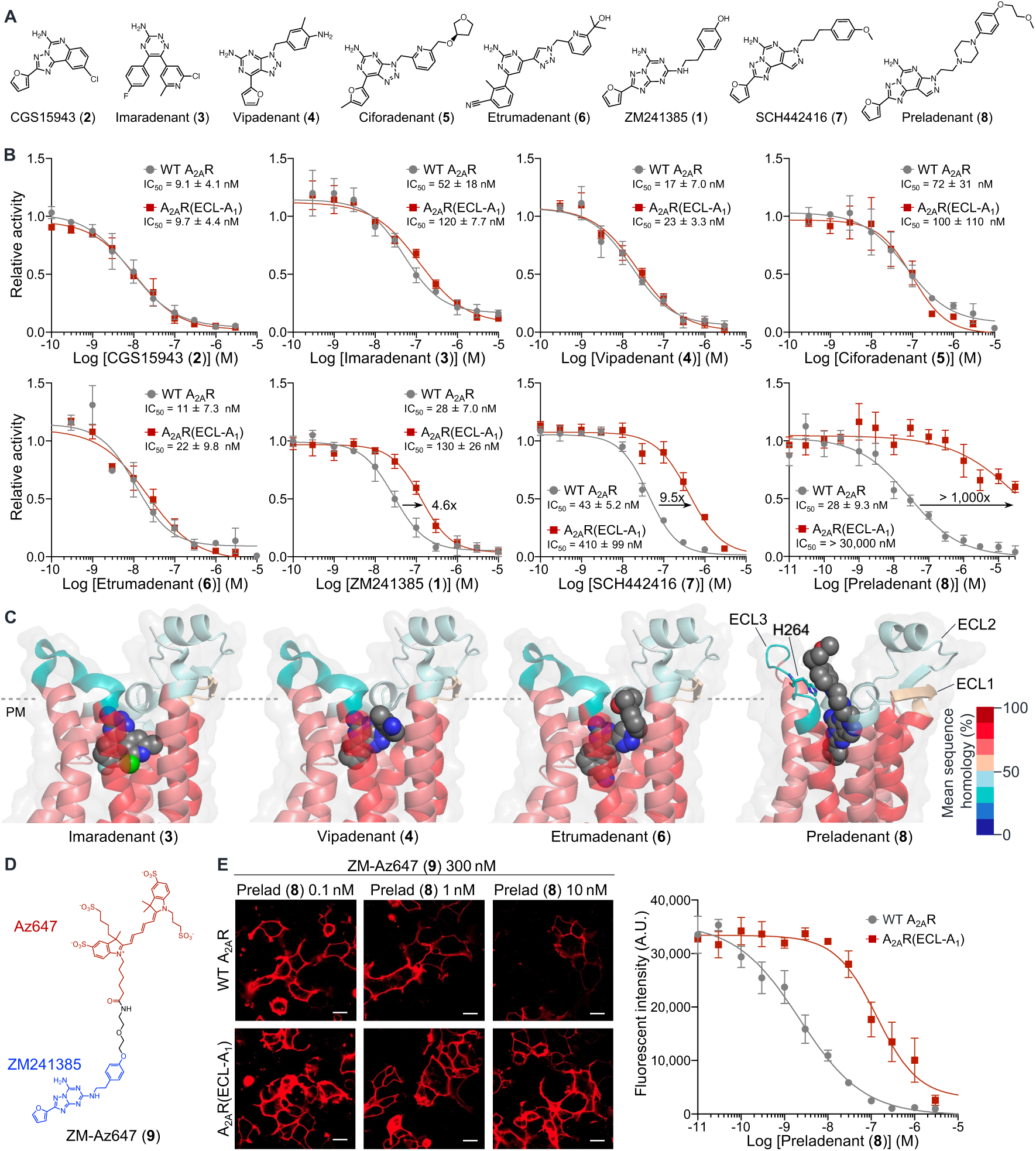
Preladenant, a suitable ligand for the antagonist-insensitive A_2A_R mutant. (**A**) Chemical structures of CGS15943 (**2**), imaradenant (**3**), vipadenant (**4**), ciforadenant (**5**), etrumadenant (**6**), ZM241385 (**1**), SCH442416 (**7**), and preladenant (**8**). (**B**) Inhibition of 100 nM Ado-induced Ca^2+^ responses by A_2A_R antagonists in HEK293 cells co-expressing Gα_15_. Concentration–response curves are shown for WT A_2A_R (gray) and A_2A_R(ECL-A_1_) (red). Relative activity is normalized to ΔRFU at 100 nM Ado. *n* = 3 or 5 independent experiments. (**C**) Binding mode of antagonists in A_2A_R. X-ray structures of A_2A_R bound to vipadenant (PDB: 5OLH), imaradenant (PDB: 6GT3), and etrumadenant (PDB: 8C9W), and estimated binding mode of preladenant determined by molecular mechanics. Receptor structures are coloured according to mean sequence homology. (**D**) Chemical structure of the fluorescent probe, ZM-Az647 (**9**). (**E**) Competitive binding assay of preladenant and ZM-Az647 to cell-surface A_2A_R. Left, representative images of HEK293 cells expressing WT A_2A_R or A_2A_R(ECL-A_1_). Scale bar, 20 µm. Right, concentration–response curves for preladenant. Cell-surface fluorescent intensity of ZM-Az647 was quantified by confocal microscopy. *n* = 3 independent experiments, 10 cells per condition. Data are presented as mean ± s.e.m.

To dissect the individual contributions of ECL2 and ECL3 mutations to preladenant (**8**) insensitivity, we generated two additional mutants: A_2A_R(ecl2-A1) and A_2A_R(ecl3-A1), in which either ECL2 or ECL3, respectively, was substituted with the corresponding A₁R sequence. The effect of each mutation on the inhibitory action of preladenant (**8**) was then evaluated. As shown in **Figure S5**, the IC_50_ values of preladenant (**8**) for A_2A_R(ecl2-A_1_) and A_2A_R(ecl3-A_1_) were shifted to higher concentrations than that of WT A_2A_R. However, these shifts were less pronounced than that observed in A_2A_R(ECL-A_1_), highlighting the importance of mutating both ECL2 and ECL3 to evade the inhibitory effect of preladenant (**8**).

We next conducted binding assays to determine whether the reduced inhibitory effect of preladenant (**8**) on A_2A_R(ECL-A_1_) is caused by a reduction in binding affinity. For this purpose, we prepared an A_2A_R-selective fluorescent probe, ZM-Az647 (**9**) (**Figure 3D**). Confocal live cell imaging showed that ZM-Az647 has high affinity for WT A_2A_R and A_2A_R(ECL-A_1_) with *K*_d_ values of 5.4 and 41 nM, respectively (**Figure S6**). Then, the affinity of preladenant (**8**) on the cell surface WT A_2A_R and A_2A_R(ECL-A_1_) were determined using a competitive assay with ZM-Az647 to derive *K*_i_ values under live cell conditions. As shown in **Figure 3E**, the fluorescence of ZM-Az647 on the cell surface decreased in the co-presence of preladenant (**8**) in a concentration-dependent manner, and *K*_i_ values for preladenant (**8**) were determined to be 0.040 and 16 nM for WT A_2A_R and A_2A_R(ECL-A_1_), respectively (**Figures 3E** and **S7**). The fold change (400-fold) was comparable to the 1,000-fold difference observed in the Ca^2+^ mobilization assay, indicating that the reduced inhibitory effect of preladenant (**8**) arises from reduced A_2A_R(ECL-A_1_) affinity to this compound.

We next evaluated other A_2A_R-dependent GPCR signaling pathways. To assess the Gα_s_ pathway under A_2A_R activation, we performed a cAMP-dependent reporter gene assay (**Figure S8A**). As shown in **Figures S8B and S8C**, adenosine-induced cAMP responses in A_2A_R(ECL-A_1_) were comparable to those observed in WT A_2A_R. Importantly, the inhibitory effect of preladenant (**8**) was significantly attenuated in A_2A_R(ECL-A_1_) compared with WT A_2A_R. We also evaluated β-arrestin signaling in WT A_2A_R and A_2A_R(ECL-A_1_). To evaluate β-arrestin translocation following GPCR activation, we employed the TANGO assay, a well-established method for monitoring this pathway (**Figure S8D**). ^21,22^ Although endogenous agonist-induced β-arrestin recruitment activities of the dopamine D_1_ receptor (D_1_R) and β_2_-adrenoceptor (β_2_AR) were robustly confirmed, the recruitment was barely detectable in either WT A_2A_R or A_2A_R(ECL-A_1_) (**Figure S8E**), indicating that A_2A_R(ECL-A_1_) does not exhibit an apparent alteration in β-arrestin signaling compared with WT A_2A_R. Overall, A_2A_R(ECL-A_1_) and preladenant (**8**) are a prominent pair for investigating the physiological roles of A_2A_R while minimizing off-target effects. Thus, we named this pair “ESCAPE-A_2A_R” (<u>e</u>xtracellular-loop <u>s</u>wapped <u>c</u>himeras <u>a</u>ctivated <u>p</u>urely by <u>e</u>ndogenous ligand for <u>A_2A_R</u>).

### Applicability of the ESCAPE strategy for other class A GPCRs

We next investigated whether the ESCAPE strategy can be extended to other class A GPCRs. Initially, we focused on the adenosine A_2B_ receptor (A_2B_R), another member of the adenosine receptor family (**Figure S9A**). Applying the ESCAPE strategy, A_2B_R(ECL-A_1_) was constructed by swapping Leu172–Val177 of ECL2 and Asn266–Lys269 of ECL3 in A_2B_R with the corresponding sequences from A_1_R (**Figure S9B**). The Ca^2+^ mobilization assay revealed that the A_2B_R(ECL-A_1_) mutant retained responsiveness to adenosine (**Figure S9C**). We then selected PSB603, a highly selective A_2B_R antagonist, for the inhibition assay (**Figure S9D**). As expected, the IC_50_ value of PSB603 for A_2B_R(ECL-A_1_) was shifted more than 170-fold higher compared with that of WT A_2B_R (**Figure S9E**). Thus, the A_2B_R(ECL-A_1_) and PSB603 pair are a suitable ESCAPE-A_2B_R system.

Next, we focused on the serotonin 2A receptor (5-HT_2A_R), which is implicated in various neuropsychiatric and neurodegenerative disorders, including schizophrenia, Alzheimer’s disease, and Parkinson’s disease.^23,24^ Although 5-HT_2A_R is a validated therapeutic target, its precise role in these disorders remains unclear, partly because several marketed antipsychotics exhibit insufficient selectivity toward 5-HT_2A_R.^25^ Therefore, elucidating the physiological roles of 5-HT_2A_R in the central nervous system (CNS) while minimizing off-target interference is crucial for understanding the mechanisms of these diseases.

To construct the ESCAPE-5HT_2A_R system, we focused on the crystal structure of 5-HT_2A_R in complex with lumateperone, a relatively selective 5-HT_2A_R antagonist approved for the treatment of schizophrenia and bipolar depression (**Figure 4, A** to **D**).^26^ This structure indicated that hydrophobic interactions between Leu228–Ala230 of ECL2 and the tetracyclic core moiety of lumateperone are critical for ligand recognition, whereas ECL3 does not participate in binding. Thus, 5-HT_2A_R(ECL-5HT_1E_) was constructed based on sequence homology by swapping Leu228–Ala230 in ECL2 of 5- HT_2A_R with the corresponding sequence of the serotonin 1E receptor (5-HT_1E_R) (**Figures 4E** and **S10**). 5-HT_2A_R(ECL-5HT_1E_) retained responsiveness to the endogenous ligand, serotonin, and the inhibitory assay showed that lumateperone antagonism was markedly reduced in the mutant (**Figures 4, F** and **G**). In detail, the IC_50_ value for 5-HT_2A_R(ECL-5HT_1E_) was shifted more than 140-fold higher compared with that of WT 5-HT_2A_R. This pair, consisting of 5-HT_2A_R(ECL-5HT_1E_) and lumateperone, can be regarded as an ESCAPE-5HT_2A_R system. Overall, the ESCAPE strategy is broadly applicable to class A GPCRs, providing a powerful platform for decoding receptor functions with high target specificity.

**Figure 4.**
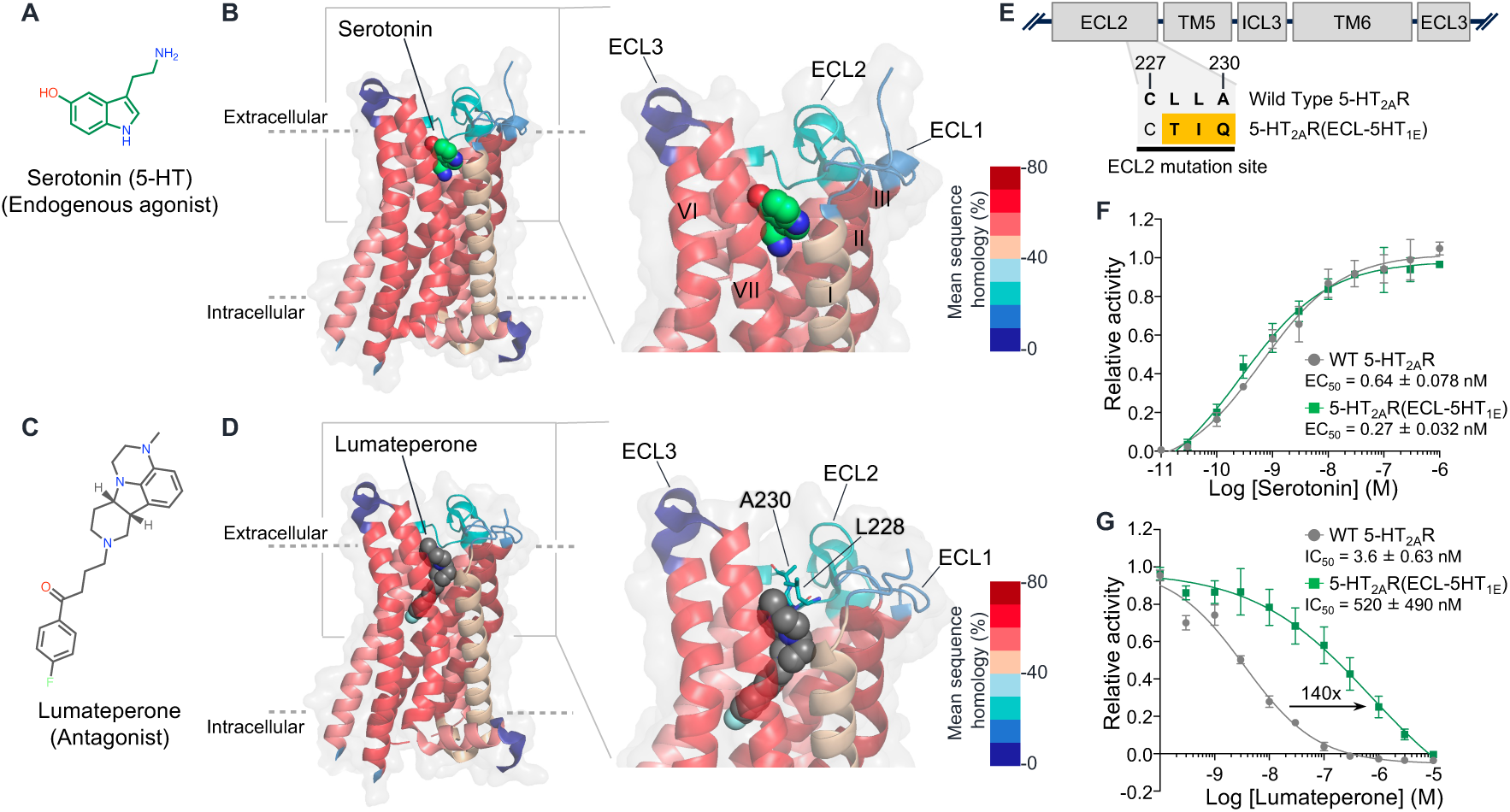
ESCAPE strategy for 5-HT_2A_R. (**A**) Chemical structure of serotonin (5-HT), the endogenous agonist of 5-HT_2A_R. (**B**) X-ray structure of 5-HT_2A_R bound to serotonin (PDB: 7WC4). Receptor structures are coloured according to mean sequence homology. (**C**) Chemical structure of lumateperone, a 5-HT_2A_R antagonist. (**D**) X-ray structure of 5-HT_2A_R bound to lumateperone (PDB: 7WC8). Receptor structures are coloured as in (B). (**E**) Mutation site of 5-HT_2A_R(ECL-5HT_1E_). Residues differing from WT 5-HT_2A_R are highlighted in yellow. (**F**) 5-HT-induced Ca^2+^ responses in HEK293 cells expressing WT 5-HT_2A_R (gray) or 5-HT_2A_R(ECL-5HT_1E_) (green). Relative activity is normalized to ΔRFU at 3 µM 5-HT. (**G**) Inhibition of 100 nM 5-HT-induced Ca^2+^ responses by lumateperone under the same expression conditions as in (F). Relative activity is normalized to ΔRFU at 100 nM 5-HT. *n* = 3 or 4 independent experiments. Data are presented as mean ± s.e.m.

### Applicability of the ESCAPE-A_2A_R system in various cell types

As described, the ESCAPE-A_2A_R system works well in HEK293 cells, a commonly used cell line model. We next tested the applicability of the ESCAPE-A_2A_R system in other cell types, in which A_2A_R functions have been previously characterized. Initially, we focused on rat pheochromocytoma (PC12) cells, which have differentiated into neuron-like cells in response to nerve growth factor (NGF) and serve as a well-established model for analyzing neuronal phenotypes (**Figure 5A**).^27^ In addition, recent studies have revealed that the expression level of A_2A_R is upregulated in brain astrocytes derived from Alzheimer’s disease patients,^13^ underscoring the importance of clarifying the roles of A_2A_R in glial cells. Therefore, the ESCAPE-A_2A_R system was also tested on primary astrocytes isolated from mouse cerebrum (**Figure 5D**).

**Figure 5.**
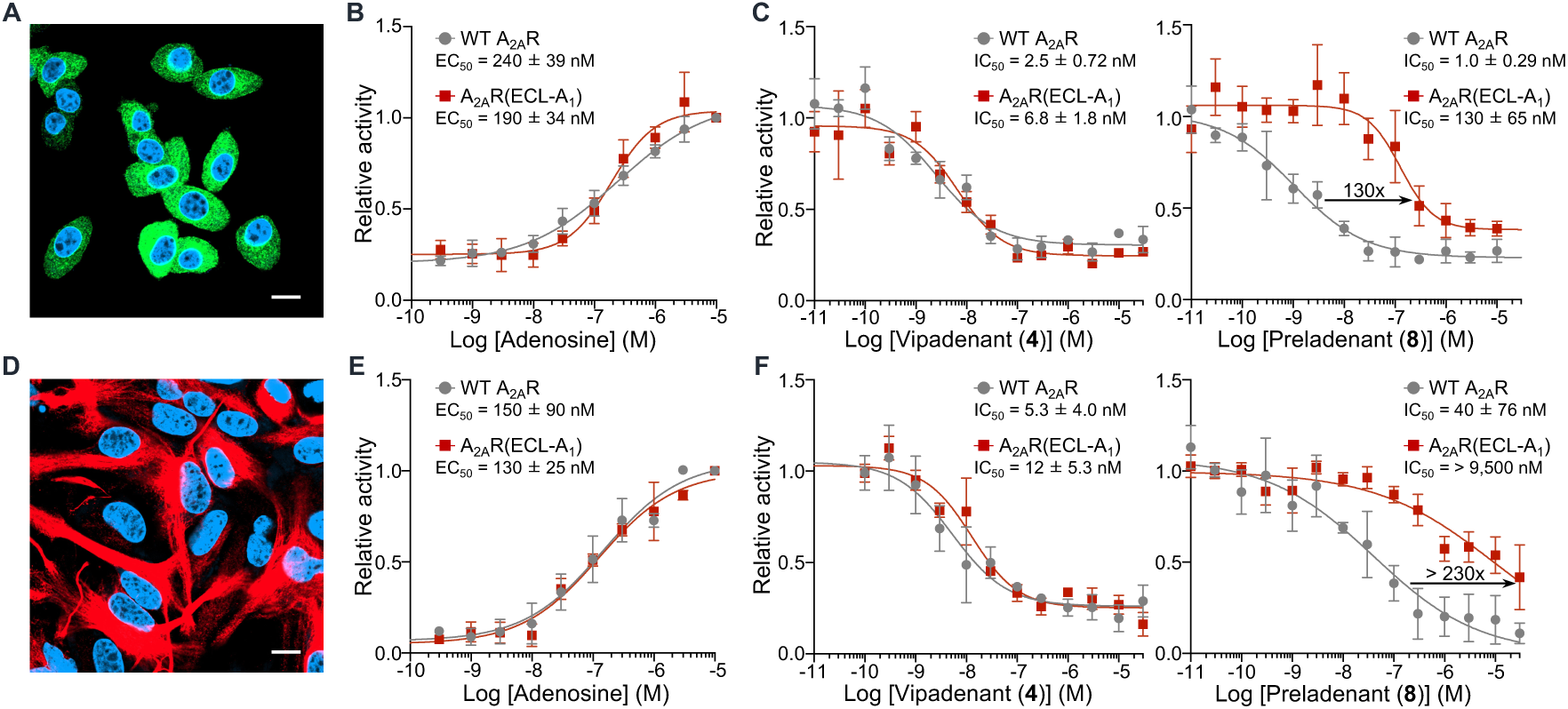
Applicability of the ESCAPE-A_2A_R system in multiple cell types. (**A**) Fluorescent visualization of PC12 cells. βIII tubulin was immunostained (green) and nuclei were counterstained with 2 µM DAPI (blue). Scale bar, 10 µm. (**B**) Ado-induced Ca^2+^ responses in PC12 cells expressed with A_2A_R (WT or ECL-A_1_) and Gα_15_. Concentration–response curves for WT A_2A_R (gray) and A_2A_R(ECL-A_1_) (red). Relative activity is normalized to ΔRFU at 10 µM Ado. (**C**) Inhibition of 1 µM Ado-induced Ca^2+^ responses by vipadenant (right) or preladenant (left) in PC12 cells. Relative activity is normalized to ΔRFU at 1 µM Ado. (**D**) Fluorescent visualization of primary cultured astrocytes. GFAP was immunostained (red) and nuclei were counterstained with 2 µM DAPI (blue). Scale bar, 10 µm. (**E**) Ado-induced Ca^2+^ responses in astrocytes expressed with A_2A_R (WT or ECL-A_1_) and Gα_15_. Experiments were performed under the same conditions as in (B). (**F**) Inhibition of 300 nM Ado-induced Ca^2+^ responses by antagonists in astrocytes. Relative activity is normalized to ΔRFU at 300 nM Ado. *n* = 3 independent experiments. Data are presented as mean ± s.e.m.

Even in PC12 cells and primary astrocytes, the adenosine-induced responses of A_2A_R(ECL-A_1_) were comparable to those of WT A_2A_R (**Figures 5, B** and **E**). In these cells, the IC_50_ values of preladenant (**8**) for A_2A_R(ECL-A_1_) were shifted to higher concentrations compared with those of WT A_2A_R (**Figures 5, C** and **F**). In contrast, such shifts were not observed with vipadenant (**4**). These results indicate that the ESCAPE-A_2A_R system is applicable to various cell types, including primary cell cultures.

### The ESCAPE systems reveal the impact of A_2A_R and 5-HT_2A_R signaling on neurite outgrowth in PC12 cells

In PC12 cells, previous studies have shown that neurite outgrowth is induced under hypoxic conditions, similar to that observed with NGF stimulation (**Figure 6A**).^28,29^ The use of an A_2A_R-targeted antagonist, CSC (8-(3-chlorostyryl)caffeine), has indicated that adenosine and A_2A_R play key roles in neurite outgrowth.^28^ However, the antagonist also inhibits monoamine oxidase B (MAO-B) with a similar affinity through a mechanism independent of A_2A_R antagonism.^30^ Additionally, hypoxic conditions promote the release of other neurotransmitters, such as dopamine (DA) and norepinephrine (NE),^31^ which may contribute to neurite outgrowth (**Figure 6A**). We applied our ESCAPE-A_2A_R system to elucidate the roles of extracellular adenosine and A_2A_R in neurite outgrowth under hypoxic conditions.

**Figure 6.**
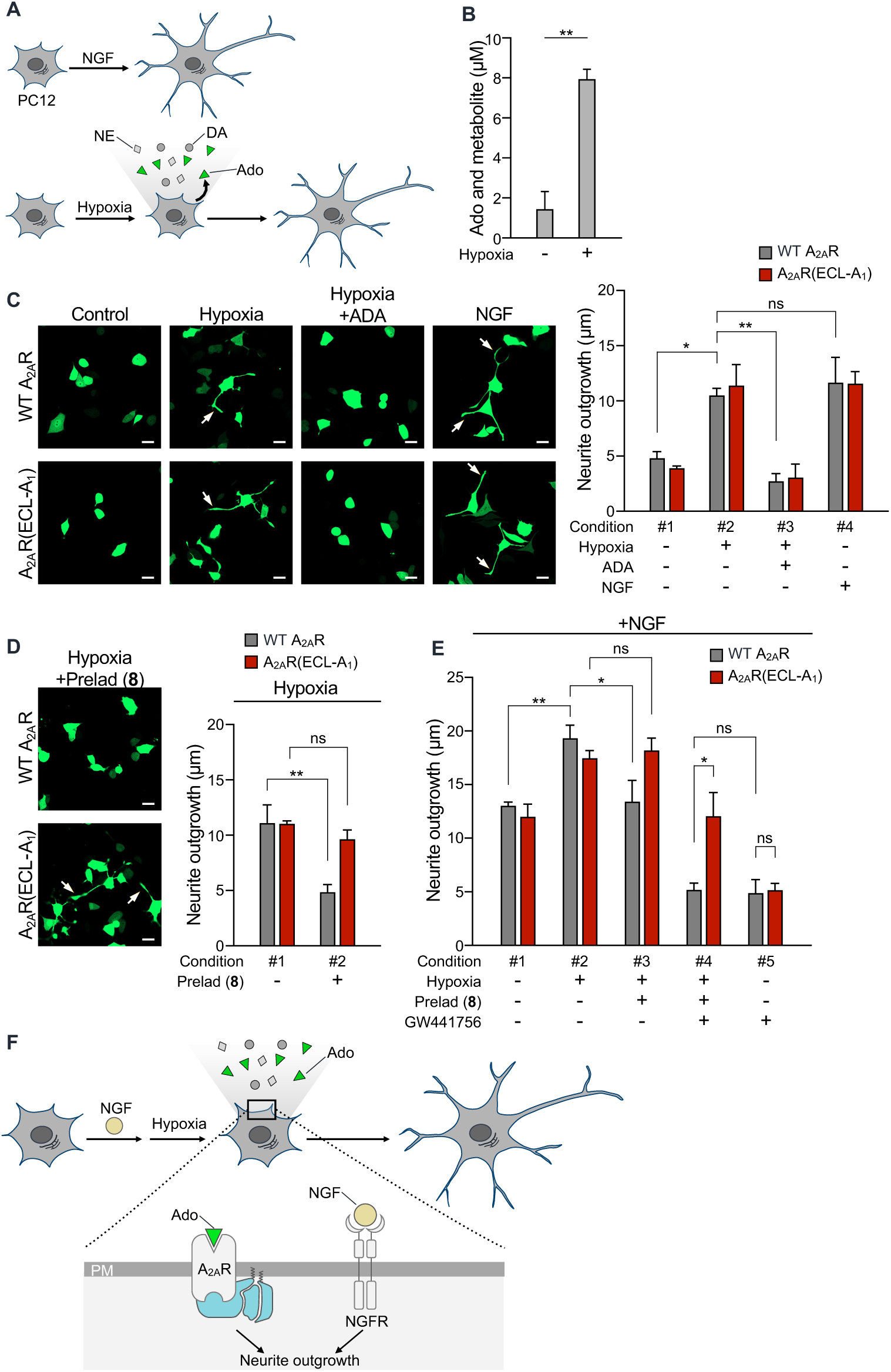
ESCAPE system revealed a critical role for A_2A_R signaling in hypoxia-induced neurite outgrowth of PC12 cells. (**A**) Schematic illustration of neurite outgrowth induced by NGF (top) and hypoxia (bottom) in PC12 cells. (**B**) Enzyme-based adenosine (Ado) detection assay. Extracellular Ado and its metabolites released from PC12 cells exposed to hypoxia (<1% O_2_) were quantified. Unpaired *t*-test. (**C** and **D**) Evaluation of hypoxia-induced Ado and A_2A_R signaling in neurite outgrowth. Left, representative images of PC12 cells expressed with A_2A_R (WT or ECL-A_1_) and AcGFP (marker). Cells in hypoxia-exposed groups were cultured under hypoxic conditions (<1% O_2_) for 12 h before imaging. Scale bar, 20 µm. Right, quantification of neurite outgrowth (longest neurite per cell). *n* = 3 independent experiments, >15 cells per condition. Two-way ANOVA with Tukey’s multiple comparisons test. (**E**) Quantification of neurite outgrowth in the presence of NGF, as in (C). (**F**) Schematic overview of the independent contributions of A_2A_R and NGF signaling to morphological changes in PC12 cells under hypoxic conditions. Data are presented as mean ± s.e.m. \**P* < 0.05, \*\**P* < 0.01; ns, not significant.

Before analyzing A_2A_R function, the extracellular adenosine released from PC12 cells under hypoxia was evaluated using an enzyme-based adenosine detection assay. As shown in **Figure 6B**, a significant increase in the concentration of released adenosine and its metabolite was detected under hypoxic conditions (< 1% O_2_) compared with normoxic conditions. Next, morphological changes were evaluated under hypoxic conditions (< 1% O_2_) in PC12 cells expressing either WT A_2A_R or A_2A_R(ECL-A_1_). As shown in **Figure 6C**, neurite outgrowth was significantly enhanced under hypoxic conditions, and the neurite length was comparable to that observed with NGF treatment (100 ng/mL) (**Figure 6C**; conditions #2 and #4, and **Figure S11A**). Importantly, treatment with adenosine deaminase (ADA; 1 U/mL), an adenosine-metabolizing enzyme, suppressed the neurite outgrowth even under the hypoxic conditions (**Figure 6C**; condition #3), suggesting that hypoxia-induced adenosine release plays a key role in promoting neurite outgrowth in PC12 cells.

We next investigated the contribution of A_2A_R signaling in neurite outgrowth under hypoxia using the ESCAPE-A_2A_R system. Under the hypoxic condition, preladenant (10 nM) treatment markedly suppressed morphological changes in WT A_2A_R-expressing cells, whereas neurite outgrowth was maintained in A_2A_R(ECL-A_1_)-expressing cells (**Figures 6D** and **S11B**). Thus, the ESCAPE-A_2A_R system showed that A_2A_R signaling plays a dominant role in hypoxia-induced neurite outgrowth in PC12 cells.

Of note, we observed that NGF-induced neurite outgrowth was further and markedly enhanced under hypoxic conditions (**Figure 6E**; conditions #1 and #2, and **Figures S11, C** and **D**). The ESCAPE-A_2A_R system was then used to evaluate whether NGF and A_2A_R signaling pathways act independently or exhibit crosstalk under hypoxic conditions. In WT A_2A_R-expressing cells under hypoxic and NGF stimuli, preladenant treatment suppressed neurite outgrowth, and the resulting neurite length was comparable to that observed with NGF-stimulation under normoxic conditions (**Figure 6E**; conditions #1 and #3). This preladenant-induced suppression was not observed in A_2A_R(ECL-A_1_)-expressing cells, indicating the importance of A_2A_R-induced signaling. Furthermore, co-treatment of preladenant (10 nM) and GW441756 (1 µM), a selective inhibitor of the NGF receptor TrkA, reduced neurite length to a level comparable to that observed under normoxic conditions in the presence of GW441756 (**Figure 6E**; conditions #4 and #5). Although the same co-treatment also suppressed neurite outgrowth in A_2A_R(ECL-A_1_)-expressing cells (**Figure 6E**; condition #4), the neurite length remained comparable to that observed under hypoxic conditions without NGF (**Figure 6D**; condition #2). Thus, our ESCAPE-A_2A_R strategy, which minimizes off-target effects, clearly indicates that adenosine signaling via A_2A_R and NGF signaling act independently and additively to promote morphological changes in PC12 cells under hypoxic conditions (**Figure 6F**).

Several studies have highlighted that serotonin receptors also play important roles in morphological changes in neuronal cells.^32,33^ We applied the ESCAPE-5HT_2A_R system to elucidate the roles of serotonin and 5-HT_2A_R in neurite outgrowth in PC12 cells. In these cells, 5-HT_2A_R(ECL-5HT_1E_) retained responsiveness to the endogenous ligand, serotonin, and antagonism by lumateperone was markedly attenuated in this mutant (**Figures S12, A** and **B**). We then examined the effects of 5-HT_2A_R signaling on the neurite outgrowth. As shown in **Figure S12C**, NGF-induced neurite outgrowth (5 ng/mL) was enhanced by in the presence of 10 nM serotonin (**Figure S12C**; conditions #1, #2 and #3). Moreover, co-treatment of 100 nM lumateperone markedly suppressed neurite outgrowth in WT 5-HT_2A_R-expressing cells, but had little effect in 5-HT_2A_R(ECL-5HT_1E_)-expressing cells (**Figure S12C**; conditions #3 and #4). Collectively, the ESCAPE-5HT_2A_R system clearly indicates that serotonin signaling through 5-HT_2A_R play an important role in NGF-induced morphological changes in PC12 cells.

## DISCUSSION

In this report, we developed a unique chemogenetic strategy, the ESCAPE-A_2A_R system, to elucidate the physiological roles of A_2A_R in living cells. The design is simple, in which ECL-swapping mutations between subtypes are sufficient to generate antagonist-insensitive mutants that retain responsiveness to adenosine. By utilizing the engineered A_2A_R in combination with preladenant (**8**), we revealed the critical role of A_2A_R in neurite outgrowth under hypoxic conditions in neuron-like cells, while minimizing off-target effects. In addition, we successfully extended the ESCAPE strategy to A_2B_R and 5-HT_2A_R, distinct members of the class A GPCR family. Given the simplicity of the ECL-swapping design, the ESCAPE strategy has a significant advantage for extending this approach to other GPCRs.

GPCRs regulate numerous cellular processes, and their dysregulation has been implicated in a wide range of diseases. Pharmacological approaches using selective inhibitors are powerful for elucidating physiological roles of target GPCRs by providing acute and reversible loss-of-function. However, even with the use of highly selective inhibitors, the potential for unintended off-target binding or the effects of metabolic byproducts cannot be fully excluded. The representative example is sitaxentan, a selective endothelin receptor A antagonist. Although this drug is used to treat pulmonary arterial hypertension, its metabolites containing a reactive quinone group can cause adverse off-target effects.^34^ Moreover, recent *in vitro* studies have uncovered previously unknown off-target activity of marketed drugs.^2,3^ The representative is lurasidone, an antipsychotic drug primarily targeting 5-HT_2A_R, which also acts on dopamine D_1_, adrenergic α_1A_, and muscarinic acetylcholine M_2_ receptors. Another example is losartan, an angiotensin II type 1 receptor blocker, which also antagonizes cannabinoid receptor 1 at nanomolar concentrations. Importantly, this interaction was overlooked during the preclinical development phase. Therefore, even when using highly selective ligands, careful evaluation of potential unforeseen off-target effects through cell-based assays is important, and we believe that the ESCAPE strategy represents a promising candidate for this purpose.

In the field of receptor chemogenetics, many powerful technologies have been developed.^35–38^ A well-established example is DREADDs, in which a designer muscarinic acetylcholine receptor is selectively activated by a synthetic ligand, such as clozapine-*N*-oxide (CNO) and the highly selective DCZ for *in vivo* applications.^37,39^ More recently, a new class of receptor chemogenetics termed PAGER has been reported, which is activated by therapeutically relevant soluble and cell-surface antigens.^40^ In these strategies, the endogenous ligand-binding properties are eliminated to enable bio-orthogonal signaling in a cell-type-specific manner. An alternative approach in chemogenetics is to understand the physiological roles of target receptors on a cell-specific basis.^41,42^ In this approach, the activity of engineered GPCRs must be regulated by a designer ligand without affecting their native function. In the current ESCAPE strategy, ECL engineering alters the affinity of selective inhibitors for target GPCRs, while preserving responsiveness to the endogenous ligand. This unique design allows the decoding of the physiological roles of individual GPCRs in living cells, free from off-target binding and the effects of metabolic byproducts.

In the ESCAPE-A_2A_R system, preladenant is used as a selective inhibitor of A_2A_R. Given that preladenant has demonstrated its efficacy in preclinical and phase I/II clinical trials with high blood–brain barrier permeability,^43^ the ESCAPE-A_2A_R system could clarify the physiological functions of A_2A_R in specific neuronal populations within the brain. In this study, both ESCAPE-A_2A_R and ESCAPE-5HT_2A_R systems were designed using X-ray structural information. Considering recent advances in structural prediction techniques such as AlphaFold3,^44^ the ESCAPE system can be applied to GPCRs whose structures remain unsolved. Moreover, designing ESCAPE platforms using approved or clinically validated antagonists will facilitate *in vivo* studies. Thus, ESCAPE-GPCR platforms may offer a valuable approach for elucidating the physiological roles of diverse GPCR subtypes *in vivo*.

## MATERIALS AND METHODS

### Synthesis

All synthesis procedures and compound characterizations are described in Supplementary information.

### Construction of expression vectors

The expression vectors pSI(A_2A_R) for A_2A_R, pSI(A_2B_R) for A_2B_R, and pCAGGS(Gα_15_) for Gα_15_ were previously described.^45^ Site-directed mutagenesis of A_2A_R was performed using the Q5^®^ Site-Directed Mutagenesis kit (NEB) according to the manufacturer’s instructions. The coding sequence of A_2A_R was subcloned into the NheI/XhoI restriction site of the pIRES2-AcGFP1 vector (Clontech) to obtain pIRES2-AcGFP1(A_2A_R) vector. 2×HA-tag sequence (<u>YPYDVPDYA</u>G<u>YPYDVPDYA</u>) was introduced at amino acid position 4 in the N-terminus region of A_2A_R using PCR-based techniques with the Q5^®^ Hot Start High-Fidelity 2X Master Mix (NEB). The resulting cDNA was subcloned into the NheI/XhoI restriction site of the pIRES2-AcGFP1 vector using the NEBuilder HiFi DNA Assembly Master Mix (NEB) to obtain pIRES2-AcGFP1(HA-A_2A_R) vector. The coding sequence of A_2A_R was subcloned into the NheI/XhoI restriction site of the pCDMd3 vector which contains CMVd3 promoter from pFN24K HaloTag CMVd3 Flexi vector (Promega) to obtain pCDMd3(A_2A_R) vector. The coding sequence of A_2A_R(ECL-A_1_) was subcloned into the EcoRI/AfeI restriction site of the A_2A_R-Tango (Addgene #66210, kindly provided by Prof. Bryan Roth)^22^ to obtain A_2A_R(ECL-A_1_)-Tango.

The cDNA region of human 5-HT_2A_R was amplified from human brain marathon-ready cDNA library (Clontech) by nested PCR method, and the coding sequence of 5-HT_2A_R was cloned into the NheI/NotI restriction site of the pSI vector (Promega) to obtain pSI(5-HT_2A_R) vector.

All NEBuilder HiFi DNA Assembly reactions were transformed into ECOS^TM^ Competent E. coli DH5α (NIPPON GENE, cat. no. 310-06236). Transformed E. coli was grown overnight at 37 °C on agar plates with 40 μg ml^−1^ kanamycin or 200 μg ml^−1^ ampicillin selection. Individual colonies were picked for liquid culture in LB media supplemented with 20 μg ml^−1^ kanamycin or 100 μg ml^−1^ ampicillin, and plasmid DNA was subsequently isolated using a NucleoSpin^®^ Plasmid Transfection-grade (TaKaRa) with endotoxin removal. All cDNA regions of the plasmids were verified by sanger sequencing (Eurofins genomics).

### Cell culture

HEK293 cells (ATCC) were cultured in a humidified atmosphere at 5% CO_2_ at 37 °C using Dulbecco’s modified Eagle’s medium (DMEM, Gibco or Nacalai) supplemented with 10% fetal bovine serum (FBS) (sigma-aldrich or Nichirei) and 100 units ml^−1^ penicillin and 100 µg ml^−1^ streptomycin.

HTLA cells (a HEK293 cell line stably expressing a tTA-dependent luciferase reporter and a β-arrestin2-TEV fusion gene) were kindly provided from the laboratory of Bryan Roth and were cultured in a humidified atmosphere at 5% CO_2_ at 37 °C using DMEM (Nacalai) supplemented with 10% FBS (Nichirei), 100 units ml^−1^ penicillin and 100 µg ml^−1^ streptomycin, 2 μg/ml puromycin and 100 μg/ml hygromycin B.

PC12 cells were cultured in a humidified atmosphere of at 5% CO_2_ at 37 °C using DMEM (Nacalai) supplemented with 10% horse serum (HS) (Gibco), 5% fetal bovine serum (FBS) (Nichirei) and 100 units ml^−1^ penicillin and 100 µg ml^−1^ streptomycin.

### Plasmid transfection in HEK293 and PC12 cells

For HEK293 cells, the culture medium was replaced with DMEM-GlutaMAX (Gibco) supplemented with 10% dialyzed FBS (Gibco or Serana Europe), 100 units ml^−1^ penicillin, and 100 µg ml^−1^ streptomycin to reduce cytotoxicity. Cells were transfected with any one group of plasmids: (a) pSI(A_2A_R) and pCAGGS(Gα_15_), (b) pSI(A_2B_R) and pCAGGS(Gα_15_), or (c) pSI(5-HT_2A_R) using Lipofectamine 3000 (Thermo Fisher Scientific) according to the manufacturer’s instructions. For co-transfections, plasmids were mixed at a 1:1 ratio by mass. Four hours after transfection, the medium was replaced with fresh medium, and the Ca^2+^ mobilization assays were performed 24 hours post-transfection.

For PC12 cells, the culture medium was replaced with DMEM supplemented with 10% HS, 5% FBS, 100 units ml^−1^ penicillin, and 100 µg ml^−1^ streptomycin. Cells were transfected with pSI(A_2A_R) and pCAGGS(Gα_15_) at 1:1 plasmid mass ratio, or with pSI(5-HT_2A_R), using Lipofectamine 3000 according to the manufacturer’s instructions. Four hours after transfection, the medium was replaced with fresh medium, and the Ca^2+^ mobilization assays were performed 24 hours post-transfection.

### Animals

All experimental procedures were performed in accordance with the National Institute of Health Guide for the Care and Use of Laboratory Animals, and were approved by the Institutional Animal Use Committees of Nagoya University. Pregnant C57BL/6N mice maintained under specific pathogen-free conditions were purchased from Japan SLC, Inc (Shizuoka, Japan). The animals were housed in a controlled environment (23 ± 1 °C, 12 h light/dark cycle) and had free access to food and water, according to the regulations of the Guidance for Proper Conduct of Animal Experiments by the Ministry of Education, Culture, Sports, Science, and Technology of Japan.

### Preparation of postnatal brain primary cell cultures

25 cm^2^ flasks (supplier) were coated with poly-D-lysine (Sigma-Aldrich) and washed with sterile dH_2_O three times. Cerebral cortices from P2–P3 C57BL/6N mice were aseptically dissected and digested with 0.05 w/v% trypsin (Fujifilm Wako) diluted in phosphate-buffered saline solution (PBS) for 20 min at 37 °C. The cells were resuspended in DMEM-GlutaMAX supplemented with 10% dialyzed FBS (Serana Europe), 100 units ml^−1^ penicillin, and 100 µg ml^−1^ streptomycin. The cells were filtered by Cell Strainer (100 µm, Falcon) followed by centrifugation at 1,000 rpm for 5 min. The cells were resuspended in DMEM-GlutaMAX supplemented with 10% dialyzed FBS, 100 units ml^−1^ penicillin, and 100 µg ml^−1^ streptomycin and plated at a density of 1.0 × 10^6^ cells in coated 25 cm^2^ flasks. The cultures were maintained in a humidified atmosphere of 95% air and 5% CO_2_ at 37 °C, changing media every 4 days.

### Electroporation of PC12 cells and primary astrocytes

Electroporation experiments were performed using the Neon™ Transfection System (Thermo Fisher Scientific) according to the manufacturer’s instructions. Briefly, PC12 cells were washed with PBS and collected using 0.05% (w/v) trypsin. Cells were resuspended in DMEM supplemented with 10% HS, 5% FBS, 100 units ml^−1^ penicillin, and 100 µg ml^−1^ streptomycin, and counted. Cells were pelleted by centrifugation at 800 rpm for 3 min, resuspended in PBS, and centrifuged again at 800 rpm for 3 min. The cell pellet was then resuspended in Buffer R (Neon™ Transfection System) at a concentration of 1.0 × 10^7^ cells ml^−1^, with 0.5 µg total DNA per 10 µl for each condition. Electroporation was carried out using 10 µl transfection tips, with three pulses at 1,300 V and a 10 ms pulse width. Cells were immediately transferred to DMEM supplemented with 1% dialyzed HS, 100 units ml^−1^ penicillin, and 100 µg ml^−1^ streptomycin, and seeded. Neurite outgrowth assays were performed 24 h post-electroporation.

For primary cultured astrocytes, cells were purified from primary cell cultures of postnatal brains on days 7–12 after plating by shaking at 250 rpm for 2 h. Cells were washed three times with PBS to remove detached cells and collected using 0.05% (w/v) trypsin. The cells were resuspended in DMEM-GlutaMAX supplemented with 10% dialyzed FBS, 100 units ml^−1^ penicillin, and 100 µg ml^−1^ streptomycin, and counted. Cells were pelleted by centrifugation at 1,000 rpm for 3 min, resuspended in PBS, and centrifuged again at 1,000 rpm for 3 min. The cell pellet was then resuspended in Buffer R at a concentration of 1.0 × 10^7^ cells ml^−1^, with 0.5 µg total DNA (pSI(A_2A_R) and pCAGGS(Gα_15_) at a 1:1 ratio by mass) per 10 µl reaction for each condition. Electroporation was performed using 10 µl transfection tips, with two pulses at 1,300 V and a 20 ms pulse width. Cells were immediately transferred to DMEM-GlutaMAX supplemented with 10% dialyzed FBS, 100 units ml^−1^ penicillin, and 100 µg ml^−1^ streptomycin, and seeded. Ca^2+^ mobilization assays were performed 24 h post-electroporation.

### Ca^2+^ mobilization assay

Transfected HEK293 cells were seeded onto 96-well advanced culture plates (Greiner Bio-One) at 2.0 × 10^4^ cells per well and incubated for 14–16 h at 37 °C under 5% CO_2_. Cells were then incubated with Cal-520 AM (final concentration 5 µM; AAT Bioquest) in DMEM-GlutaMAX supplemented with 10% dialyzed FBS, 100 units ml^−1^ penicillin, and 100 µg ml^−1^ streptomycin for 4 h at 37 °C under 5% CO_2_.

For PC12 cells, transfected cells were seeded onto 96-well advanced culture plates at 4.0 × 10^4^ cells per well and incubated for 14–16 h at 37 °C under 5% CO_2_. Cells were washed with DMEM supplemented with 5% FBS, 100 units ml^−1^ penicillin, and 100 µg ml^−1^ streptomycin. The culture medium was then replaced with DMEM containing 10 µM Cal-520 AM, 2.5 mM probenecid (an inhibitor of organic anion transporters), 5% FBS, 100 units ml^−1^ penicillin, and 100 µg ml^−1^ streptomycin, and cells were incubated for 4 h at 37 °C under 5% CO_2_.

For primary cultured astrocytes, electroporated cells were seeded onto 96-well advanced TC plates at 2.5 × 10^4^ cells per well and incubated for 14–16 h at 37 °C under 5% CO_2_. The culture medium was then replaced with DMEM-GlutaMAX containing 10 µM Cal-520 AM, 10% dialyzed FBS, 100 units ml^−1^ penicillin, and 100 µg ml^−1^ streptomycin, and cells were incubated for 4 h at 37 °C under 5% CO_2_.

Test compounds were diluted to the indicated concentrations in pre-warmed HEPES-buffered saline (HBS; 20 mM HEPES pH 7.4, 107 mM NaCl, 6 mM KCl, 1.2 mM MgSO_4_, 2 mM CaCl_2_, 11.5 mM glucose) and kept at 37 °C. Cells were washed three times with HBS using an AquaMax 2000 (Molecular Devices). Cells were stimulated with test compounds, and the fluorescence intensity was measured at 525 nm (excitation at 490 nm) using a FlexStation 3 (Molecular Devices). The changes in fluorescence were fitted with GraphPad Prism (version 10) with a four-parameter logistic equation: a + (b-a)/(1 + (x/c)^d). The half-maximal effective concentration (EC_50_) and half-maximal inhibitory concentration (IC_50_) values were calculated.

### Confocal live cell imaging of cell-surface A_2A_Rs in HEK293 cells

HEK293 cells were transfected with pIRES2-AcGFP1(HA-A_2A_R) or pIRES2-AcGFP1(A_2A_R) vectors using Lipofectamine 3000 according to the manufacturer’s instructions. Twenty-four hours after transfection, cells were seeded onto poly-L-lysine-coated glass-bottom dishes and incubated for an additional 24 h at 37 °C under 5% CO_2_. To investigate the expression levels of A_2A_R mutants on the plasma membrane by immunostaining, cells were washed with HBS and incubated with Alexa Fluor^®^ 647-conjugated anti-HA tag antibody (Cell Signaling Technology, #37297; 1:200 dilution in HBS) for 60 min at room temperature to prevent endocytosis and then washed twice with HBS prior to imaging.

For binding assays on the cell surface A_2A_R, cells were washed twice with HBS and incubated with ZM-Az647 (an A_2A_R-selective fluorescent probe) diluted in HBS to the indicated concentrations. For competitive inhibition assays, cells expressing A_2A_R were incubated with 300 nM ZM-Az647 and indicated concentrations of preladenant for 30 min at room temperature to allow binding to reach equilibrium prior to imaging.

Confocal live imaging was performed with LSM900 confocal microscope (Carl Zeiss) equipped with a 63× oil-immersion objective (numerical aperture (NA) 1.4). Fluorescence images were acquired by excitation at 488 or 640 nm derived from diode lasers. Cell surface fluorescent intensity was quantified by line scan analysis of transfection marker-positive cells by ZEN blue software (Carl Zeiss) and calculated with subtraction of background. The changes in fluorescent intensity were fitted with GraphPad Prism (version 10) using a four-parameter logistic equation: a + (b-a)/(1 + (x/c)^d). The dissociation constant (*K*_d_) values of ZM-Az647 and half-maximal inhibitory concentration (IC_50_) values of preladenant were calculated. The inhibition constant (*K*_i_) values for preladenant were calculated from the IC₅₀ values using the Cheng-Prusoff equation: *K*_i_ = IC_50_/(1 + ([*L*]/*K*_d_)), where [*L*] is the concentration of ZM-Az647 (300 nM).

### Molecular mechanics calculation

The structure of A_2A_R in complex with preladenant was modeled based on the structure of A_2A_R in complex with PSB-2113 (a preladenant conjugate), which was determined by X-ray crystallography (PDB ID: 7PX4).^46^ First, PSB-2113 was replaced by preladenant by Avogadro software^47,48^, keeping the common atoms at the same positions. Second, the redundant N-terminal residues of A_2A_R were deleted, and the ICL3 was modeled using the comparative function of Modeller^49^ implemented in ChimeraX^50^. The single point mutation (S91^3.39^K) introduced in the crystal structure was maintained. Amber ff19SB^51^ and GAFF2^52,53^ force fields were employed for A_2A_R and preladenant, respectively. After performing the partial charge calculation based on the AM1-BCC model^54,55^ using the Antechamber module in AmberTools23^48^, the force field parameter and coordinate files were generated by using the tLEaP module in AmberTools23^53^. Finally, we performed energy minimization in vacuo using the Sander module in AmberTools23^53^ to obtain the model structure of the A_2A_R/preladenant complex.

### cAMP-dependent luciferase reporter gene assay

96-well opaque plates (Nunc) were coated with 0.01% collagen solution (Fujifilm Wako) and washed with PBS. HEK293 cells were transfected with pCDM-d3(A_2A_R) and pGL4.29(Luc2P-CRE) (Promega) vectors using Lipofectamine 3000 according to the manufacturer’s instructions. Transfected cells were seeded onto collagen-coated 96-well plates at 2.0 × 10^4^ cells per well and incubated for 14–16 h at 37 °C under 5% CO_2_. Test compounds were diluted to the indicated concentrations in pre-warmed DMEM- GlutaMAX supplemented with 10% dialyzed FBS, 100 units ml^−1^ penicillin, and 100 µg ml^−1^ streptomycin and kept at 37 °C. Cells were stimulated with test compounds, and incubated for 4 h at 37 °C under 5% CO_2_. 100 μl per well of Bright-Glo solution (Promega) was added to each well. After incubation for 3-5 min at room temperature, luminescence was counted using a FlexStation 3 (Molecular Devices). Relative luminescence units (RLU) were exported into Excel spreadsheets, and GraphPad Prism was used for analysis of data.

### β-arrestin-recruitment assay

96-well opaque plates (Nunc) were coated with 0.01% collagen solution (Fujifilm Wako) and washed with PBS. HTLA cells were transfected with A_2A_R-Tango (Addgene # 66210), A_2A_R(ECL-A_1_)-Tango, D_1_R-Tango (Addgene # 66268) and β_2_AR-Tango (Addgene # 66220) vectors using Lipofectamine 3000 according to the manufacturer’s instructions. Transfected cells were seeded onto collagen-coated 96-well plates at 2.0 × 10^4^ cells per well and incubated for 14–16 h at 37 °C under 5% CO_2_.

Test compounds were diluted to the indicated concentrations in pre-warmed DMEM-GlutaMAX supplemented with 10% dialyzed FBS, 100 units ml^−1^ penicillin, and 100 µg ml^−1^ streptomycin and kept at 37 °C. Cells were stimulated with test compounds, and incubated for 8 h at 37 °C under 5% CO_2_. 100 μl per well of Bright-Glo solution (Promega) was added to each well. After incubation for 3-5 min at room temperature, luminescence was counted using a FlexStation 3 (Molecular Devices). Relative luminescence units (RLU) were exported into Excel spreadsheets, and GraphPad Prism was used for analysis of data.

### Immunostaining of PC12 cells and primary astrocytes

PC12 cells were washed with PBS and fixed with 4% paraformaldehyde (PFA) for 15 min. After three washes with PBS, fixed cells were permeabilized with 0.1% Triton X-100 for 10 min and washed three times again with PBS. Blocking was performed using Blocking One Histo (Nacalai Tesque) for 10 min. Following a PBS wash, cells were incubated with anti- β III tubulin antibody (abcam, ab18207) in PBS containing 5% Blocking One Histo (1:1,000 dilution) for 1 h at room temperature. After three washes with PBS, cells were incubated with Alexa Fluor^®^ 488-conjugated anti-rabbit IgG antibody (Cell Signaling Technology, #4412s) in PBS containing 5% Blocking One Histo (1:1,000 dilution) for 1 h at room temperature. After another three washes with PBS, cells were incubated with 2 µM 4’,6-diamidino-2-phenylindole (DAPI) for 5 min.

For primary cultured astrocytes, the same protocol was followed except that the primary antibody was anti-GFAP antibody (Pharmingen, 60311D; 1:1,000 dilution), and the secondary antibody was Alexa Fluor^®^ 647-conjugated anti-mouse IgG antibody (Invitrogen, A-21235; 1:1,000 dilution). The secondary antibody incubation was carried out for 24 h at 4 °C.

Imaging was performed with LSM900 confocal microscopy. Fluorescence images were acquired by excitation at 405, 488 or 640 nm derived from diode lasers.

### Adenosine detection assay in cultured medium of PC12 cells

PC12 cells were seeded onto poly-L-lysine-coated 4-well plate (SPL Life Sciences) at 1.0 × 10^5^ cells per well in 350 µl medium and incubated for 16–20 h at 37 °C under 5% CO_2_. For hypoxic treatment, plate was placed in CO_2_ incubator with Anaero Pack (Mitsubishi Gas Chemical) to maintain <1% O_2_. Normoxic samples were incubated in parallel. After 24 h, 200 µl culture supernatants were collected and centrifuged at 10,000 × rpm for 5 min at 4 °C to remove insoluble particles. Concentration of adenosine and its metabolites was measured using the Adenosine Assay Kit MET-5090 (Cell Biolabs) according to the manufacturer’s instructions with minor modifications. Briefly, 50 μl of each supernatant sample was added to two wells of a 96-well black plate (Corning). One well was treated 50 μl of Reaction Mix (containing adenosine deaminase, purine nucleoside phosphorylase, and xanthine oxidase), while the other was treated Control Mix without enzymes to determine background fluorescence.

### Evaluation of neurite outgrowth in PC12 cells

For morphological analysis under A_2A_R or 5-HT_2A_R expression conditions, electroporated PC12 cells (A_2A_R) or transfected PC12 cells (5-HT_2A_R) were seeded onto poly-L-lysine-coated glass-bottom dishes at 0.5 × 10^5^ cells per dish and incubated for 12 h at 37 °C under 5% CO_2_. Cells were treated with test compounds diluted in DMEM supplemented with 1% dialyzed HS, 100 units ml^−1^ penicillin, and 100 µg ml^−1^ streptomycin. For hypoxic treatment, dishes were placed in CO_2_ incubator with an Anaero Pack to maintain <1% O_2_ for 12 h. Normoxic samples were incubated in parallel.

Imaging was performed with an LSM900 confocal microscope. Fluorescence images were acquired by excitation at 488 nm derived from diode lasers. Neurite outgrowth was quantified by measuring the longest neurite per cell using ImageJ.

### Statistical analysis

All data are presented as mean ± s.e.m. We accumulated the data for each condition from at least three independent experiments. We evaluated statistical significance with unpaired *t*-test, one-way ANOVA with Dunnett’s test, or two-way ANOVA with Tukey’s multiple comparisons test. A value of *P* < 0.05 was considered significant.

## Supporting information

Supplementary Informations

## ACKNOWLEDGMENTS

We thank Mr. Takumitsu Shakuno (Nagoya University) for the construction of plasmids, Mr. Takuto Sugawara (Nagoya University) for the synthesis of the fluorescent probe precursor, Mr. Shuntaro Kashiwa (Nagoya University) for technical support, and Dr. Andrew Dingley from Edanz (https://jp.edanz.com/ac) for editing a draft of this manuscript. We thank Prof. Yasuo Mori (Kyoto University) for kindly providing PC12 cells. We also thank Prof. Bryan Roth (University of North Carolina) for kindly providing HTLA cells and Tango plasmids. This work was funded by Grants-in-Aid for Scientific Research (KAKENHI) (Grant Number 24KJ1275 to Y.M., 22K05351 to T.D., 23K14154 to D.P.T., 23H02445, 23H02424, 23H04058, 24H01357, and 24H02259, to A.K., 24H02265, 24H00492, and 24K21823 to S.K.), Novartis Foundation (to S.K.), the Takeda Science Foundation (to S.K.) and supported AMED Grant Number 24zf0127012 to S.K.

## AUTHOR CONTRIBUTIONS

Y.M., T.D., and S.K. initiated and designed the project. Y.M. performed the construction and characterization of the ESCAPE-A_2A_R and the ESCAPE-A_2B_R. Y.M. and H.I. performed the construction and characterization of the ESCAPE-5-HT_2A_R. Y.M. performed synthesis, adenosine detection assay, and confocal imaging. T.H. and D.P.T. performed molecular mechanics calculations. Y.M. and S.K. wrote the manuscript. T.D. and A.K. reviewed and edited the manuscript. All authors discussed and commented on the manuscript.

