## Supplementary Informations for "Off-target-free chemogenetic platform that decodes physiological roles of target GPCRs"

**This PDF file includes:**

Supplementary Methods

Figures S1 to S12

Supplementary Reference

### Supplementary Methods

#### General materials and methods for organic synthesis

All chemical reagents and solvents were purchased from commercial suppliers (Tokyo Chemical Industry (TCI), Fujifilm Wako Pure Chemical, Kanto Chemical, Sigma-Aldrich, or Fluoroprobes) and used without further purification. Solvents and additives are abbreviated as follows: DMF = *N*,*N*-dimethylformamide, DIPEA = *N*,*N*-diisopropylethylamine, TFA = trifluoroacetic acid, NHS = N-hydroxysuccinimide. Thin layer chromatography (TLC) was performed on silica gel 60 $F_{254}$ precoated glass plates (Merck Millipore) and visualized via fluorescence quenching and ninhydrin staining. Chromatographic purification was performed using Wakosil C-200 (spherical, 64–210 μm, Fujifilm Wako) and silica gel 60 N (spherical, neutral, 40–50 μm, Kanto Chemical). Matrix-assisted laser desorption/ionization time-of-flight mass spectroscopy (MALDI-TOF MS) spectra were recorded on an autoflex maX instrument (Bruker Daltonics) using dithranol (DIT) as the matrix. High-resolution mass spectra (HRMS) were measured on a compact (Bruker) equipped with electron spray ionization (ESI). Reversed-phase HPLC (RP-HPLC) was performed on a Hitachi Chromaster system equipped with a 5410 UV detector, a 5110 pump, and a YMC-Pack ODS-A column (S-5 μm, 250 × 20 mm). UV/Vis detection was at 250 nm. All runs used linear gradients of $CH_3CN$ (solvent A) and 10 mM ammonium acetate aqueous solution (solvent B).

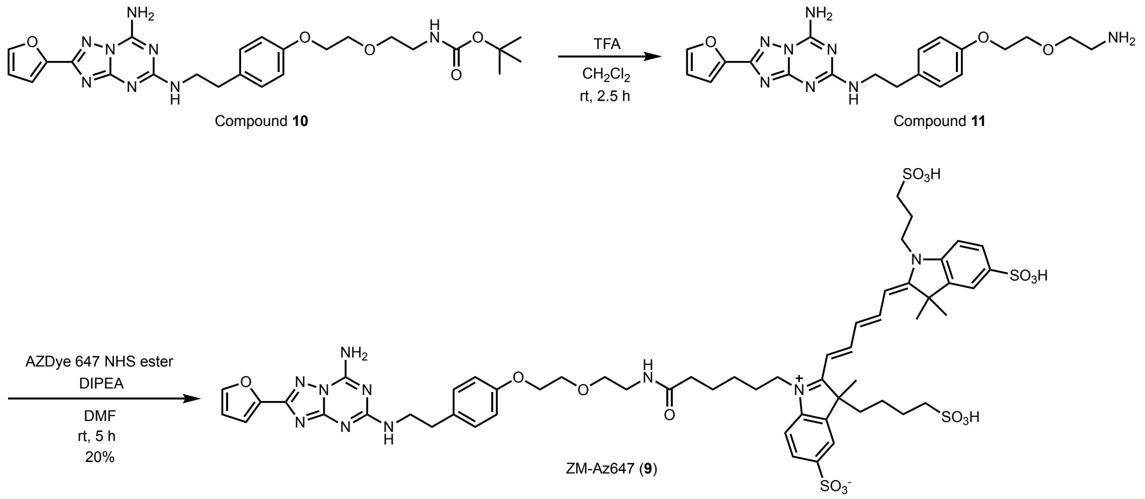


**Scheme. Synthesis of ZM-Az647 (9).**

Compound **10** were synthesized as previously reported[S1].

ZM-Az647 (**9**)

TFA (0.60 ml, 894 mg, 7.8 mmol) was added to a solution of compound **10** (18 mg, 34.3 μmol) in dry $CH_2Cl_2$ (2.4 ml). The reaction mixture was stirred for 2.5 h at room temperature. After confirming the consumption of compound **10**, the reaction mixture was combined with toluene and concentrated *in vacuo* to obtain compound **11**. HRMS (ESI) *m*/*z* calcd for $C_{20}H_{25}N_8O_3^+$ $[M+H]^+$ 425.2044, found 425.2017. A portion of compound **11** was used without further purification in the following reaction. Compound **11** (2.06 μmol, estimated on the assumption of quantitative Boc deprotection) and AZDye 647 NHS ester (1.0 mg, 1.03 μmol) were dissolved in dry DMF (392 μl). To this solution, DIPEA (11.7 μl, 69 μmol) was added under $N_2$. The mixture was stirred at room temperature for 5 h. The crude mixture was purified by RP-HPLC (ODS-A, 250 × 10 mm, mobile phase; solvent A/solvent B = 5 : 95 to 30 : 70 (linear gradient over 40 min), flow rate; 3 ml/min, detection; UV (250 nm)) to give **ZM-Az647** (0.58 μmol was determined based on the molecular extinction coefficient of AZDye 647 NHS ester (270,000 $M^{-1}cm^{-1}$), 56% yield) as a blue solid. HRMS (ESI) *m*/*z* calcd for $C_{57}H_{67}N_{10}O_{16}S_4^{3-}$ $[M-3H]^{3-}$ 425.1212 found 425.1253.

A

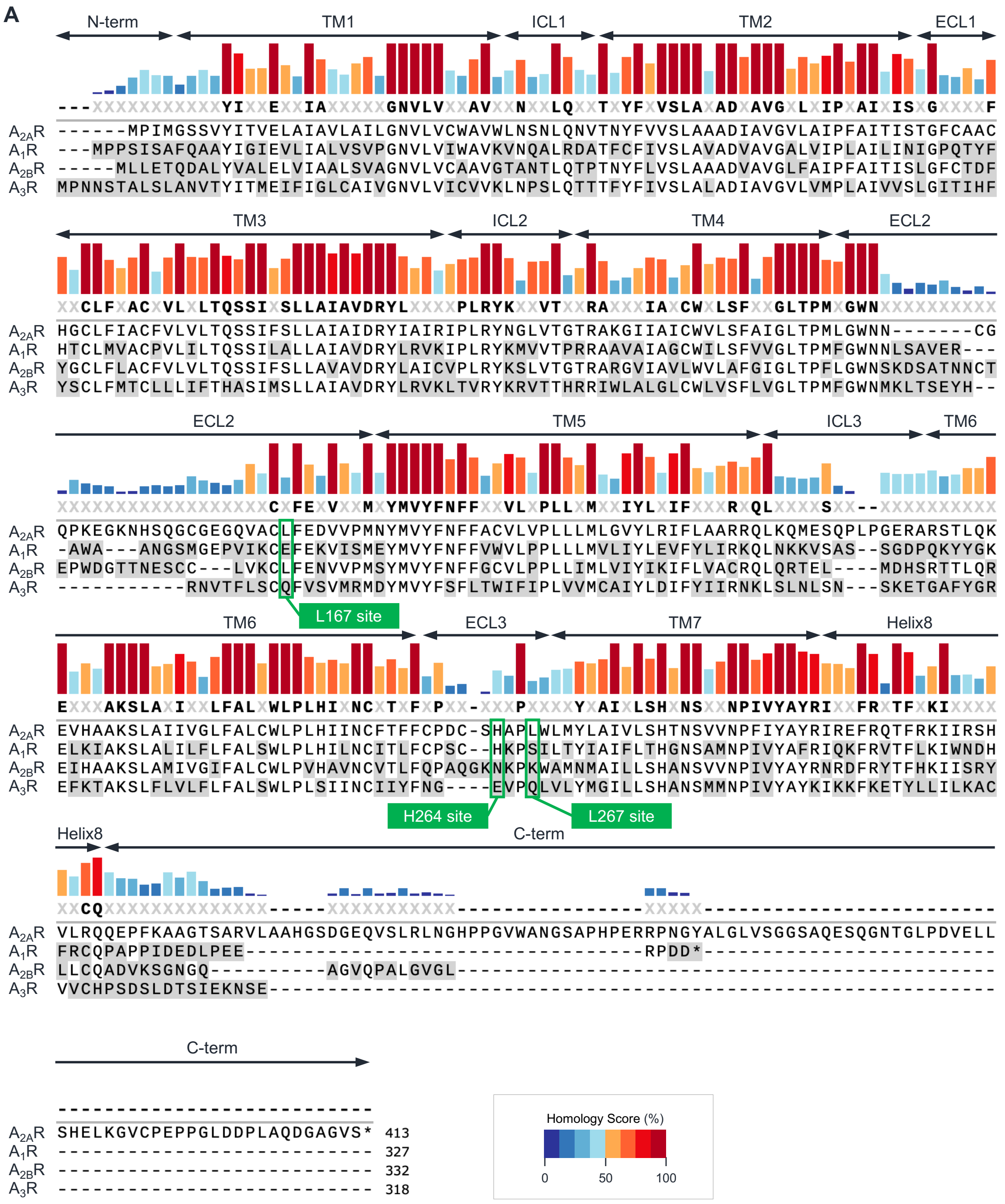


B

| Domain Name | N-term | TM1 | ICL1 | TM2 | ECL1 | TM3 | ICL2 | TM4 | ECL2 | TM5 | ICL3 | TM6 | ECL3 | TM7 | Helix8 | C-term |
|---|---|---|---|---|---|---|---|---|---|---|---|---|---|---|---|---|
| Mean sequence homology (%) | 38.5 | 65.1 | 55.8 | 81.9 | 52.4 | 80.1 | 69.5 | 69.5 | 44.9 | 72.3 | 33.9 | 73.3 | 36.3 | 77.8 | 59.7 | - |

**Figure S1. Sequence homology of human adenosine receptor subtypes**

(**A**) Multiple-sequence alignment of the four human adenosine receptor subtypes generated using ClustalW.

(**B**) Mean sequence homology within each structural domain of the $A_{2A}R$. Domain boundaries were defined according to the previous report[15].

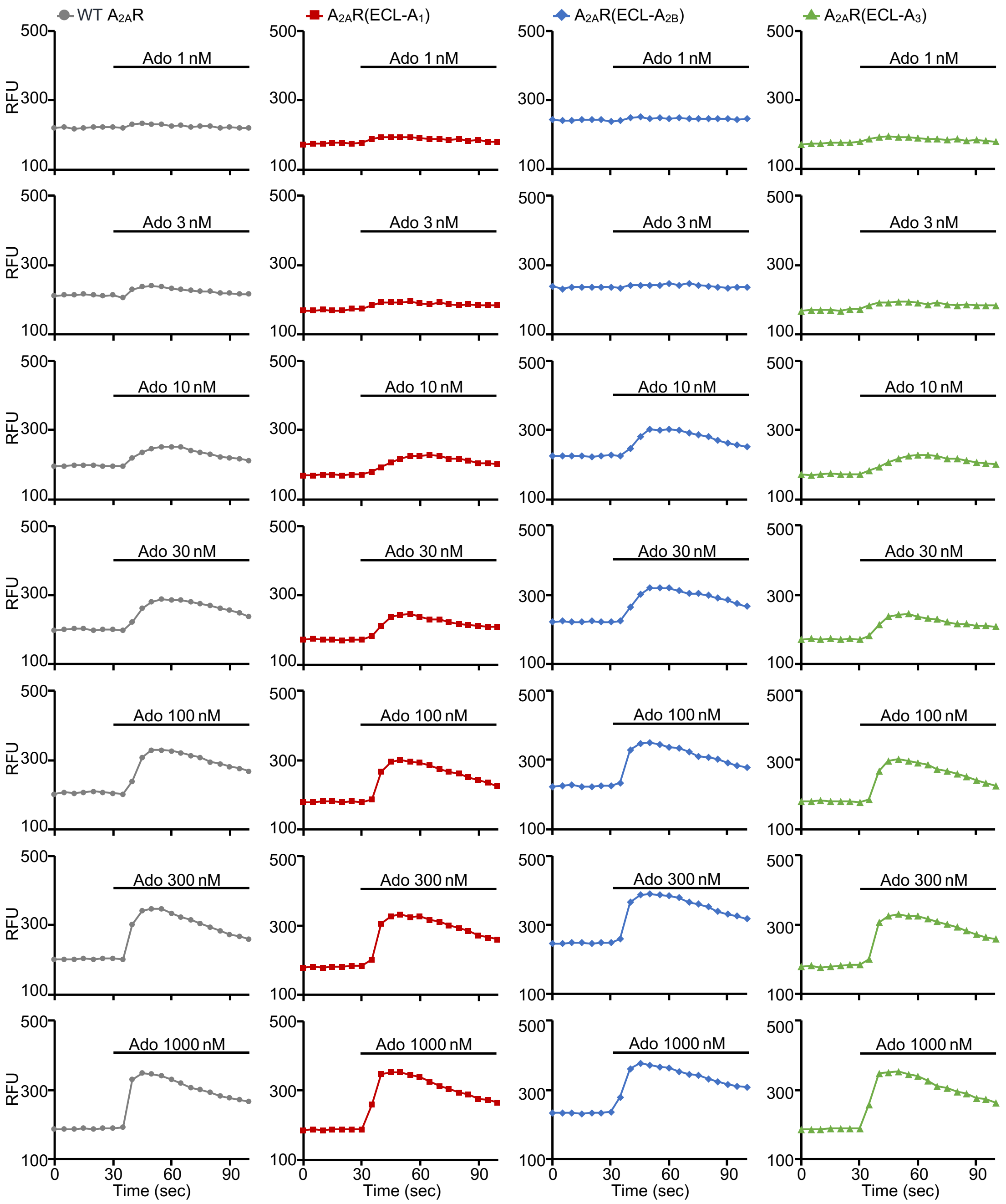


**Figure S2. Representative traces of adenosine (Ado)-induced $Ca^{2+}$ responses**

Cal-520 fluorescence signals in HEK293 cells co-expressing $G\alpha_{15}$ and either WT $A_{2A}R$ (gray), $A_{2A}R$(ECL-$A_1$) (red), $A_{2A}R$(ECL-$A_{2B}$) (blue), or $A_{2A}R$(ECL-$A_3$) (green), stimulated by Ado (black bar). Fluorescence signals are shown as relative fluorescence units (RFU).

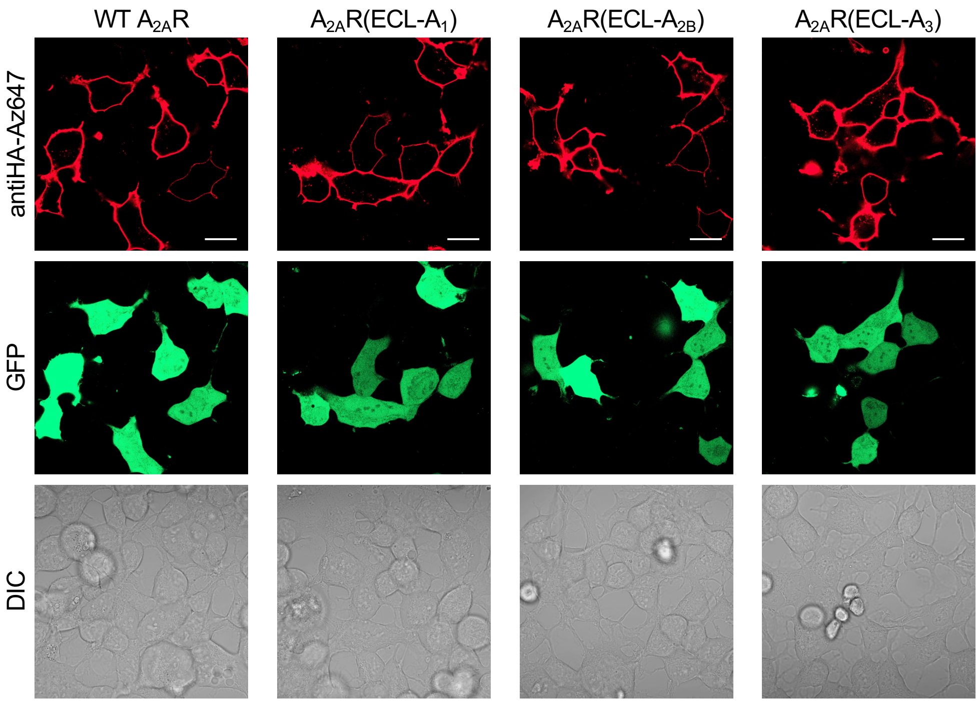


**Figure S3. Cell-surface expression of WT $A_{2A}R$ and chimeric $A_{2A}R$ mutants**

Representative images of HEK293 cells expressing 2×HA-tagged WT $A_{2A}R$ or $A_{2A}R$ mutants. Fluorescence signals from Alexa 647-conjugated anti-HA tag antibody (top), GFP (middle), and differential interference contrast (DIC; bottom) are shown. Scale bar, 20 μm.

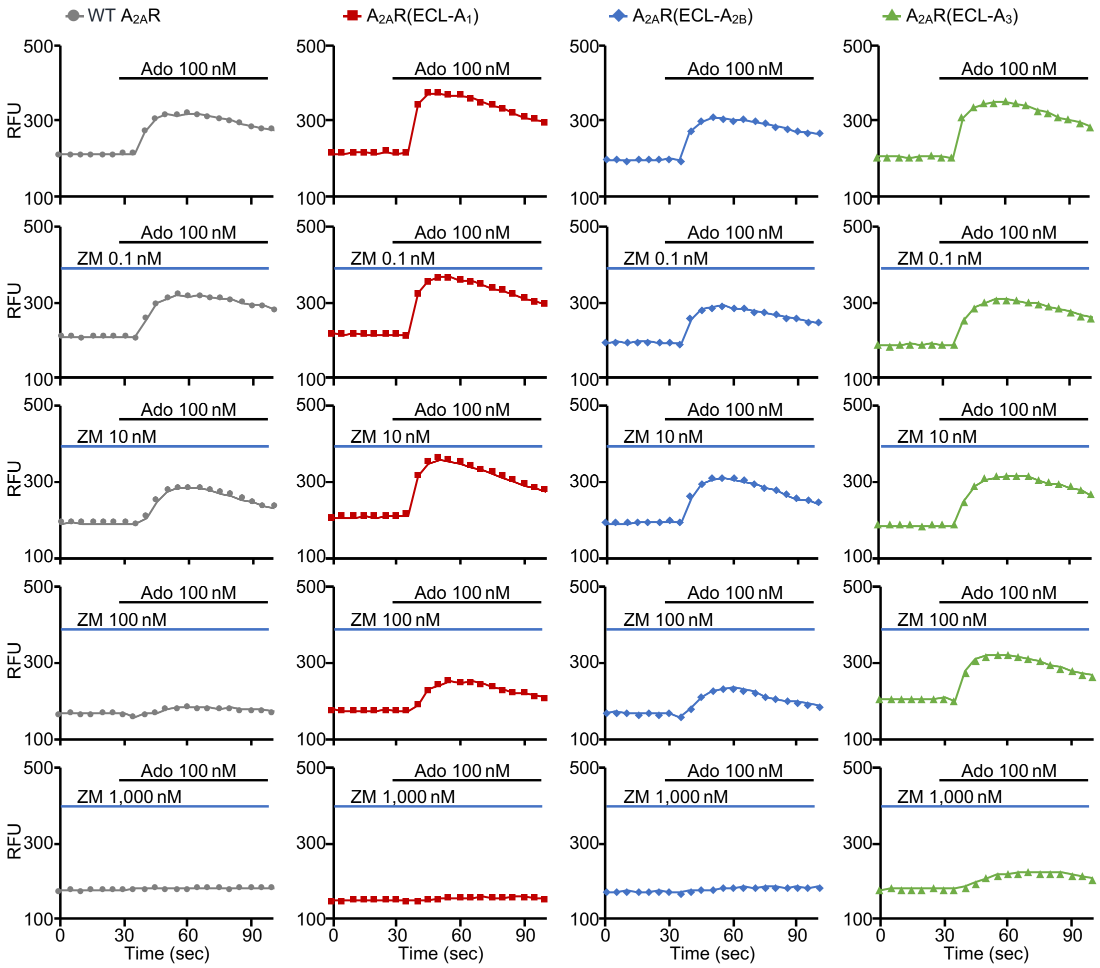


**Figure S4. Representative traces of Ado-induced $Ca^{2+}$ responses inhibited by ZM241385**

Cal-520 fluorescence signals in HEK293 cells co-expressing $G\alpha_{15}$ and either WT $A_{2A}R$ (gray), $A_{2A}R$(ECL-$A_1$) (red), $A_{2A}R$(ECL-$A_{2B}$) (blue), or $A_{2A}R$(ECL-$A_3$) (green). Cells were pre-treated with ZM241385 (ZM, blue bar) followed by stimulated by 100 nM Ado (black bar). Fluorescence signals are shown as relative fluorescence units (RFU).

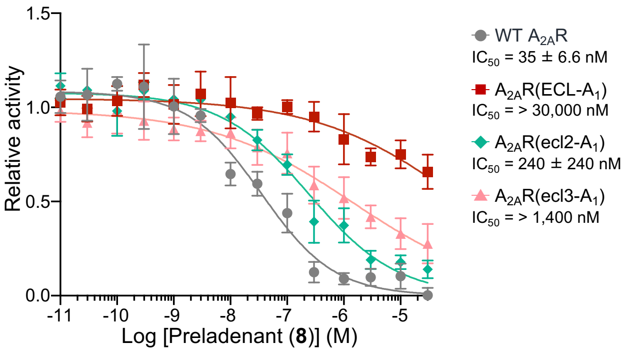


**Figure S5. Individual contributions of ECL2 and ECL3 mutations to preladenant insensitivity**

Inhibition of 100 nM Ado-induced $Ca^{2+}$ responses by preladenant in HEK293 cells co-expressing $G\alpha_{15}$ and either WT $A_{2A}R$ (gray), $A_{2A}R$(ECL-$A_1$) (red), $A_{2A}R$(ecl2-$A_1$) (light-green), or $A_{2A}R$(ecl3-$A_1$) (pink). Relative activity is normalized to ΔRFU at 100 nM Ado. *n* = 3 independent experiments. Data are presented as mean ± s.e.m.

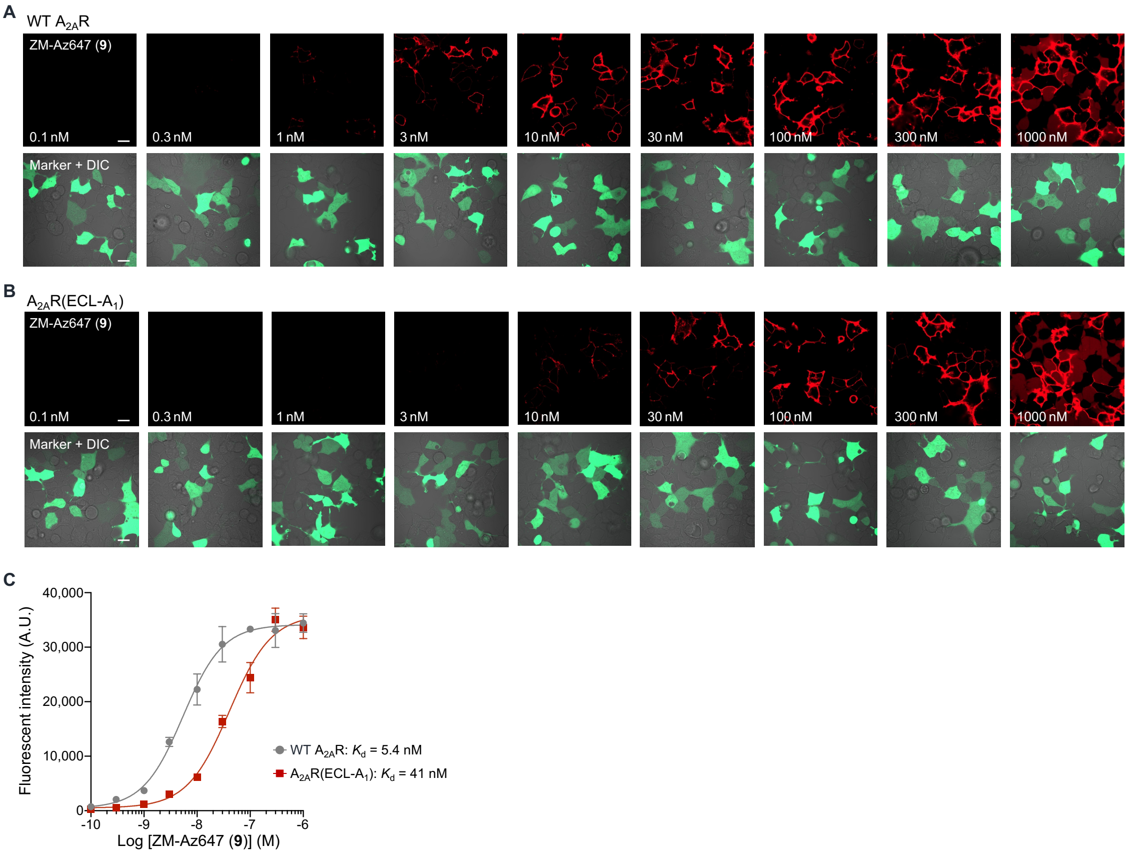


**Figure S6. Cell-surface binding of ZM-Az647 to $A_{2A}R$**

(**A** and **B**) Representative images of HEK293 cells expressing WT $A_{2A}R$ (A) and $A_{2A}R(ECL\text{-}A_1)$ (B), stained with fluorescent probe ZM-Az647 (red signal). AcGFP indicates transfected cells (green signal). Scale bar, 20 µm.

(**C**) Concentration–response curves for ZM-Az647. Cell-surface fluorescent intensity was quantified by confocal microscopy. $n = 3$ independent experiments, 10 cells per condition. Data are presented as mean ± s.e.m.

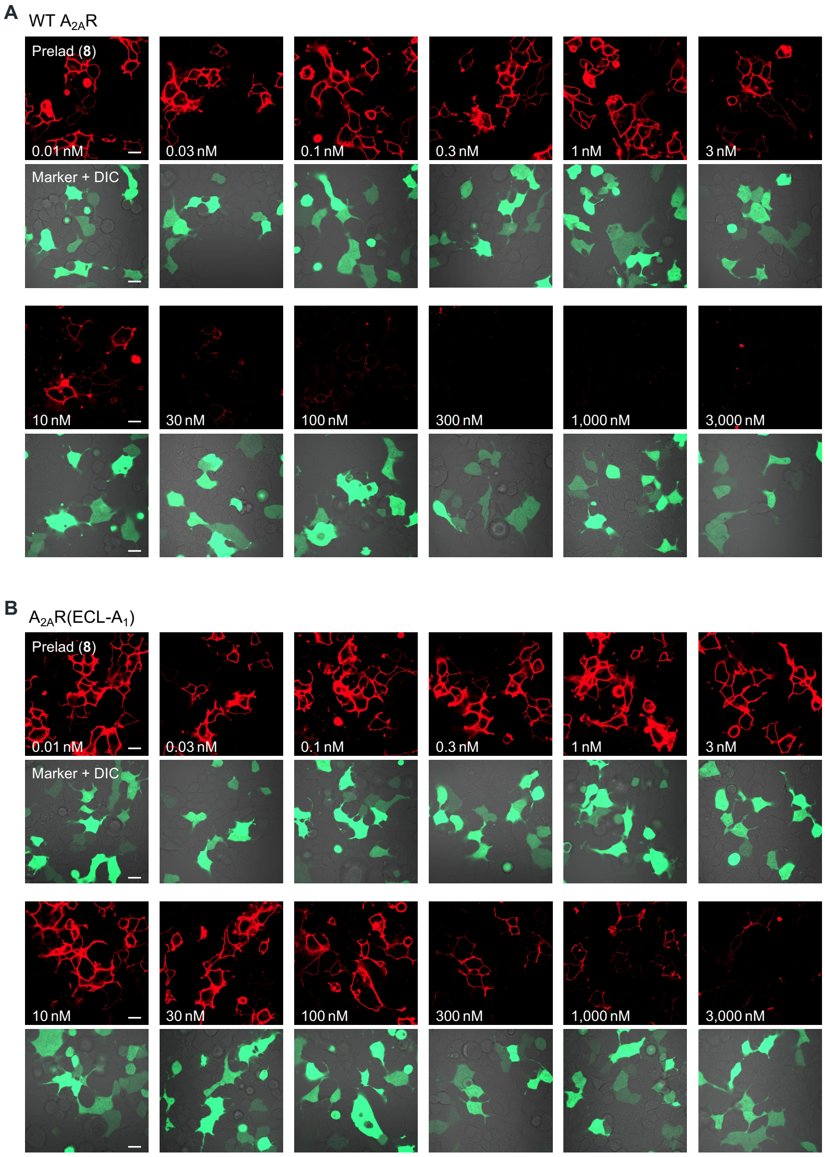


**Figure S7. Competitive binding assay of preladenant and ZM-Az647 to cell-surface $A_{2A}R$**

(**A** and **B**) Representative images of HEK293 cells expressing WT $A_{2A}R$ (A) and $A_{2A}R(ECL\text{-}A_1)$ (B), stained with ZM-Az647 (red signal) in the presence of preladenant. AcGFP indicates transfected cells (green signal). Scale bar, 20 µm.

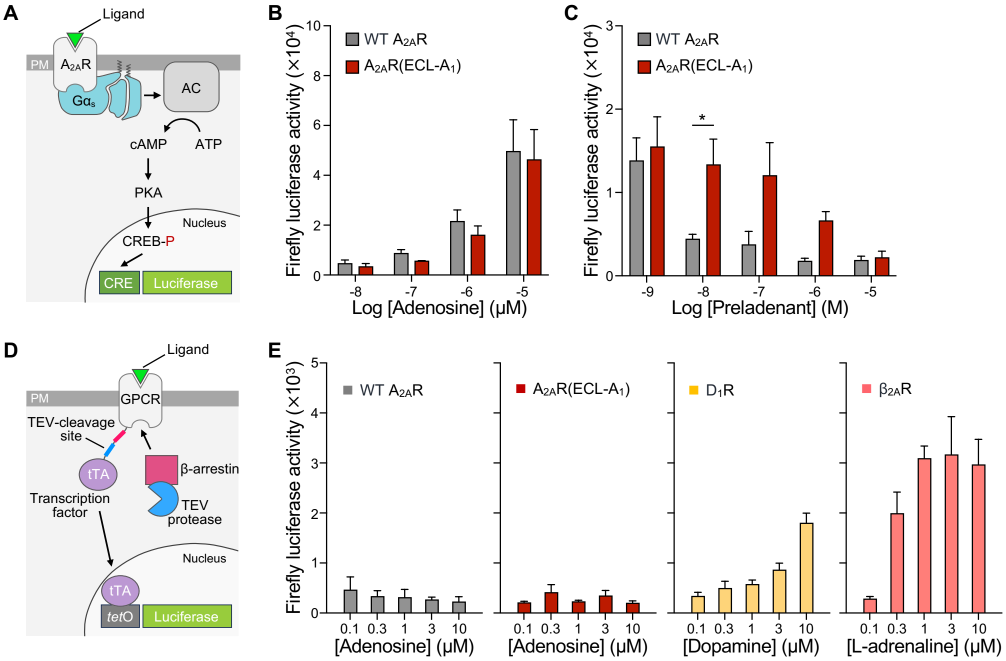


**Figure S8. Evaluation of G$\alpha_s$ and β-arrestin pathway under $A_{2A}R$ activation**

(**A**) Schematic illustration of cAMP-dependent reporter gene assay.

(**B**) Ado-induced cAMP responses in HEK293 cells co-expressing CRE-luc2P and either WT $A_{2A}R$ (gray) or $A_{2A}R(ECL\text{-}A_1)$ (red).

(**C**) Inhibition of 1,000 nM Ado-induced cAMP responses by Preladenant under the same expression conditions as in (B). Two-way ANOVA with Šídák's multiple comparisons test. *$P < 0.05$.

(**D**) Schematic illustration of β-arrestin translocation following GPCR activation (Tango) assay.

(**E**) Endogenous agonist-induced β-arrestin recruitment activities of WT $A_{2A}R$ (gray), $A_{2A}R(ECL\text{-}A_1)$ (red), the dopamine $D_1$ receptor ($D_1R$) (yellow), and $\beta_2$-adrenoceptor ($\beta_{2A}R$) (orange). $N$ = 3-5 independent experiments. Data are presented as mean ± s.e.m.

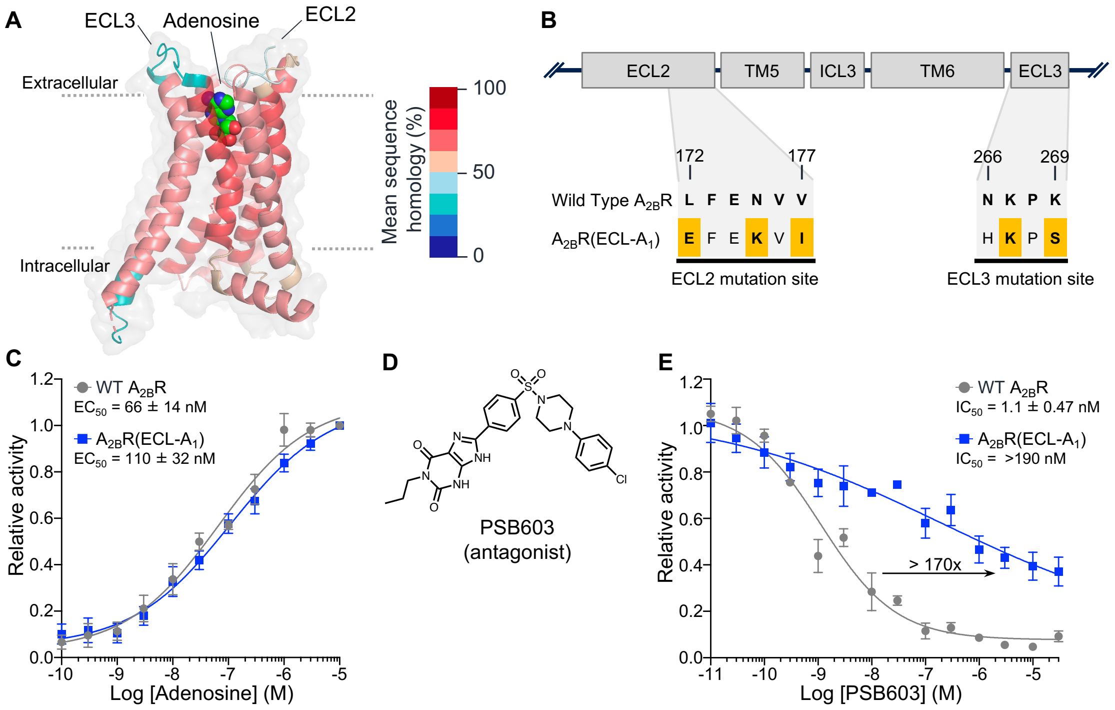


**Figure S9. Construction and characterization of ESCAPE-$A_{2B}R$**

(**A**) Cryo-EM structure of $A_{2B}R$ bound to Ado (PDB: 8HDP). Receptor structure of $A_{2B}R$ is coloured according to mean sequence homology.

(**B**) Mutation sites of the $A_{2B}R(ECL\text{-}A_1)$. Residues differing from WT $A_{2B}R$ are highlighted in yellow.

(**C**) Ado-induced $Ca^{2+}$ responses in HEK293 cells co-expressing $G\alpha_{15}$ and either WT $A_{2B}R$ (gray) or $A_{2B}R(ECL\text{-}A_1)$ (blue). Relative activity is normalized to ΔRFU at 10 µM Ado.

(**D**) Chemical structure of PSB603, an $A_{2B}R$-selective antagonist.

(**E**) Inhibition of 300 nM Ado-induced $Ca^{2+}$ responses by PSB603 under the same expression conditions as in (C). Relative activity is normalized to ΔRFU at 300 nM Ado. $n$ = 3 independent experiments. Data are presented as mean ± s.e.m.

**A**

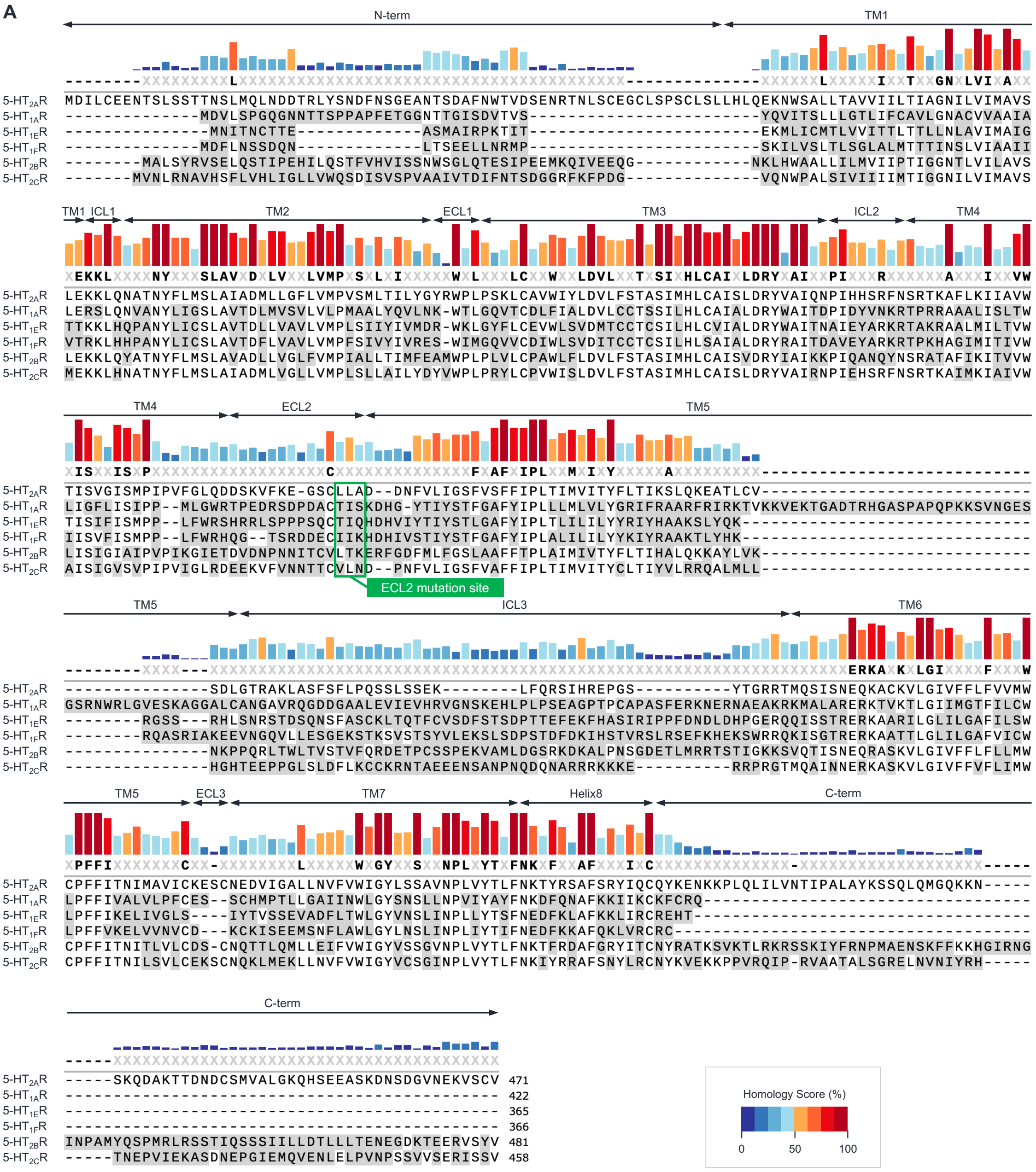


**B**

| Domain name | N-term | TM1 | ICL1 | TM2 | ECL1 | TM3 | ICL2 | TM4 | ECL2 | TM5 | ICL3 | TM6 | ECL3 | TM7 | Helix8 | C-term |
|---|---|---|---|---|---|---|---|---|---|---|---|---|---|---|---|---|
| Mean sequence homology (%) | 15.8 | 50.0 | 50.0 | 72.4 | 18.8 | 70.6 | 62.5 | 57.6 | 29.6 | 52.8 | 20.0 | 64.7 | 8.3 | 63.6 | 54.5 | 9.1 |

**Figure S10. Sequence homology of human 5-$HT_{2A}$ and 5-HT receptor subtypes**

(**A**) Multiple-sequence alignment of human 5-HT receptor subtypes generated using ClustalW.

(**B**) Mean sequence homology within each structural domain of 5-$HT_{2A}$R compared with 5-$HT_{1E}$R, calculated using Clustal Omega.

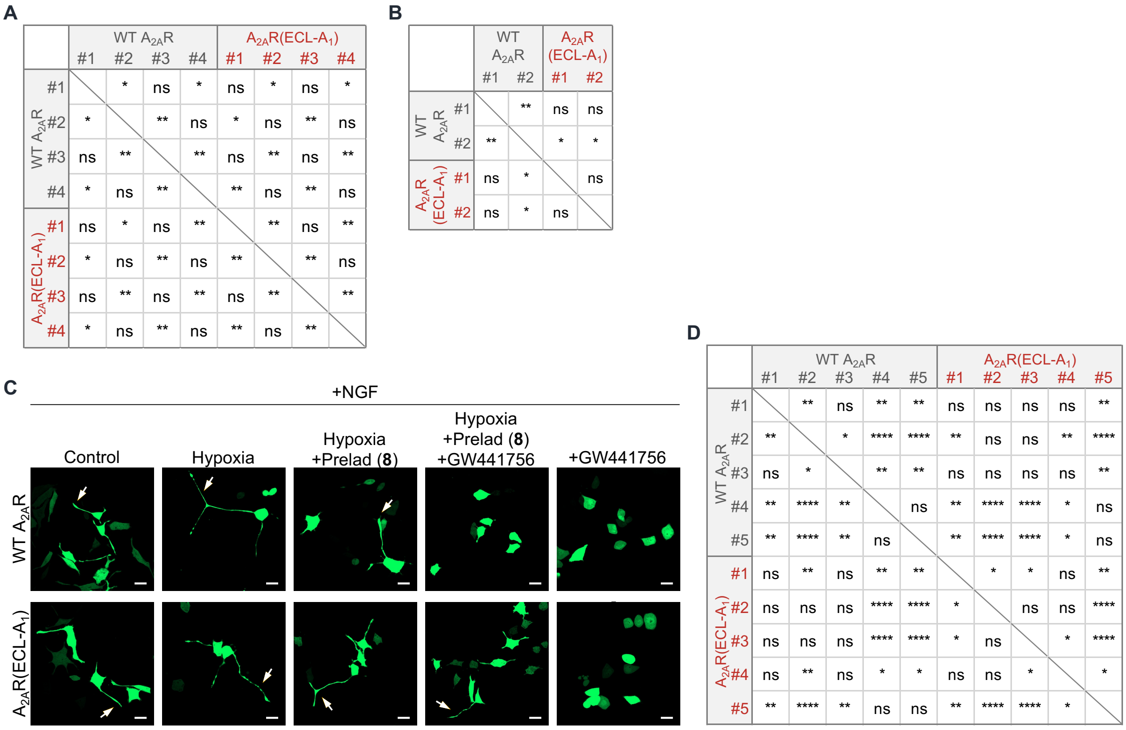


**Figure S11. Evaluation of hypoxia-induced Ado–$A_{2A}R$ signaling and NGF signaling in neurite outgrowth**

(**A** and **B**) Statistical analyses in **Figure 5C** (A) and **Figure 5D** (B) using two-way ANOVA with Tukey's multiple comparisons test.

(**C**) Evaluation of neurite outgrowth in the presence of NGF. Representative images of PC12 cells expressed with $A_{2A}R$ (WT or ECL-$A_1$) and AcGFP (marker) are shown. Cells in hypoxia-exposed groups were cultured under hypoxic conditions (<1% $O_2$) for 12 h before imaging. Scale bar, 20 μm.

(**D**) Statistical analysis in **Figure 5E**, as in (A). *$P < 0.05$, **$P < 0.01$, ***$P < 0.001$, ****$P < 0.0001$; ns, not significant.

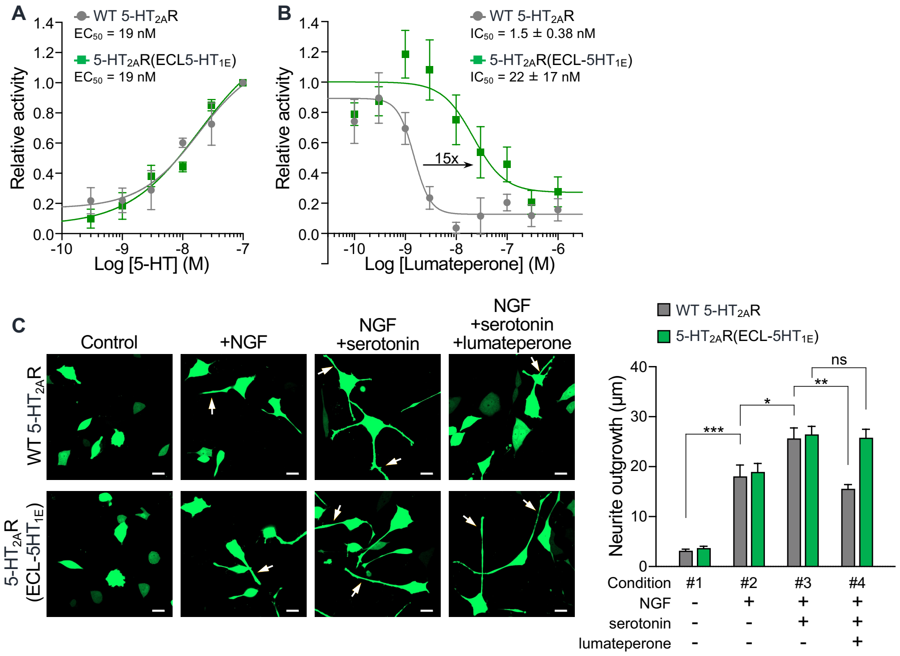


**Figure S12. Evaluation of 5-HT$_{2A}$R signaling in neurite outgrowth**

(**A**) 5-HT-induced $Ca^{2+}$ responses in PC12 cells expressed with WT 5-HT$_{2A}$R (gray) or 5-HT$_{2A}$R (ECL-5HT$_{1E}$) (green). Relative activity is normalized to ΔRFU at 100 nM 5-HT.
(**B**) Inhibition of 100 nM 5-HT-induced $Ca^{2+}$ responses by lumateperone under the same expression conditions as in (A). Relative activity is normalized to ΔRFU at 100 nM 5-HT. $n$ = 3-4 independent experiments. Data are presented as mean ± s.e.m.
(**C**) Evaluation of 5-HT-induced 5-HT$_{2A}$R signaling in neurite outgrowth. Left, representative images of PC12 cells expressed with 5-HT$_{2A}$R (WT or ECL-5HT$_{1E}$) and AcGFP (marker). Scale bar, 20 μm. Right, quantification of neurite outgrowth (longest neurite per cell). $n$ = 3 independent experiments, >30 cells per condition. Two-way ANOVA with Tukey's multiple comparisons test. *$P < 0.05$, **$P < 0.01$, ***$P < 0.001$; ns, not significant.

S1. Fukasawa, G., Matsuoka, Y., Tran, D.P., Nishigaki, H., Fukunaga, K., Watanabe, T., Doura, T., Terasaka, N., Kagawa, A., Murata, T., et al. (2026). *In Vitro* Evolution of the Adenosine $A_{2A}$ Receptor Based on an Antagonist Binding Using a Ribosome Display. J. Am. Chem. Soc. 148, 11415–11423. https://doi.org/10.1021/jacs.6c02372.
